# Prefrontal representations can specialize rather than generalize with experience

**DOI:** 10.64898/2026.03.27.714523

**Authors:** Uladzislau Barayeu, Andrea Cumpelik, Karola Kaefer, Jozsef Csicsvari

## Abstract

While the medial prefrontal cortex (mPFC) interacts with the hippocampus to support learning and consolidation, how its neural representations evolve remains unclear. We recorded population activity from CA1 and the mPFC as rats trained on radial maze tasks over weeks. Low-dimensional mPFC manifolds generated using UMAP or PCA embeddings initially tracked session progression, showing within-session drift. As animals gained task familiarity, this temporal tracking weakened; mPFC manifolds differentiated specific arms, mirroring hippocampal representations. Arm discriminability was selectively enhanced at locations requiring spatial choices, but weakened when such demands were absent. These findings demonstrate that systems consolidation does not uniformly promote cortical generalization. Instead, the mPFC may slowly refine representations in response to specific task demands to support tailored task schemas.

## Introduction

The hippocampus and medial prefrontal cortex (mPFC) coordinate their activity to support spatial memory, contextual learning, and goal-directed navigation (Euston et al., 2012; Ito et al., 2015; Spellman et al., 2015; Kitamura et al., 2017). Their changing role has been studied in relation to spatial or contextual learning when recall occurred several days or weeks after task acquisition (Frankland et al., 2004; Maviel et al., 2004; Tonegawa et al., 2018). Neuronal recordings indicate that mPFC populations typically represent abstract information related to spatial and contextual coding (Morrissey et al., 2017; Yu et al., 2018; Kaefer et al., 2020; Samborska et al., 2022; Tang et al., 2023; El-Gaby et al., 2024; Muysers et al., 2025). Furthermore, behavioral experiments suggest that remote memories undergo generalization over time, characterized by reduced contextual specificity or a reliance on egocentric rather than allocentric reference frames (Packard and McGaugh, 1996; Wiltgen and Silva, 2007; Wiltgen and Tanaka, 2013; Morrissey et al., 2017). These findings are consistent with consolidation theories proposing that abstract, gist-like representations develop in the mPFC over repeated experience (Nadel and Moscovitch, 1997; Frankland and Bontempi, 2005; Robin and Moscovitch, 2017; Lutz et al., 2026). Computational neuroscience has also incorporated these ideas into the Complementary Learning Systems (CLS) model (McClelland et al., 1995; Schapiro et al., 2017), which is also central to modern machine learning approaches to lifelong learning (Kumaran et al., 2016). The CLS model assumes that the neocortex extracts statistical invariants to build generalized knowledge. This abstraction is also consistent with experimental data linking the mPFC to behavioral schema formation (Tse et al., 2007a, 2011a; McKenzie et al., 2014a; Farzanfar et al., 2023a).

Often, behavior requires the differentiation of spatial features even after consolidation has occurred. How cortical schemas maintain task specificity in such cases remains unknown. Here, we investigated the dynamics of representational change in the mPFC as rats trained on radial maze tasks and acquired proficiency over a two-week period. We reasoned that the mPFC undergoes a slow, experience-dependent reorganization of its population geometry. By tracking population activity along low-dimensional manifolds, we found that experience drove a geometric shift from temporal tracking to task-specific spatial coding, demonstrating that cortical consolidation is dynamically optimized by behavioral utility.

## Results

We recorded from CA1 and medial prefrontal cortex (mPFC) ensembles in two separate cohorts of rats, each trained on one of two radial eight-arm maze tasks with distinct cognitive requirements (Figure 1). For the spatial reference and working memory (SRW) task, we used two subcohorts of rats (n = 4 each); the first cohort received a dual implant consisting of a 24-tetrode microdrive for CA1 and a Neuropixels probe for mPFC. In the second cohort, mPFC Neuropixels recordings spanned initial learning and subsequent task repetition days to assess the long-term stabilization of representations. These animals collected rewards from three fixed arms, reaching near-optimal performance within 3 to 4 days (Figure 1B, Figure S1). In the cue-guided association (CGA) task, the identity of a food cue presented in the start arm indicated which of the two arms was baited. This task was significantly more difficult, requiring at least 8 days of training with up to 70 trials per day to reach proficiency (Figure 1A, Figure S1A,B). In these experiments, rats were implanted with a 32-tetrode microdrive targeting each region with 16 tetrodes (n=3).

**Figure 1:**
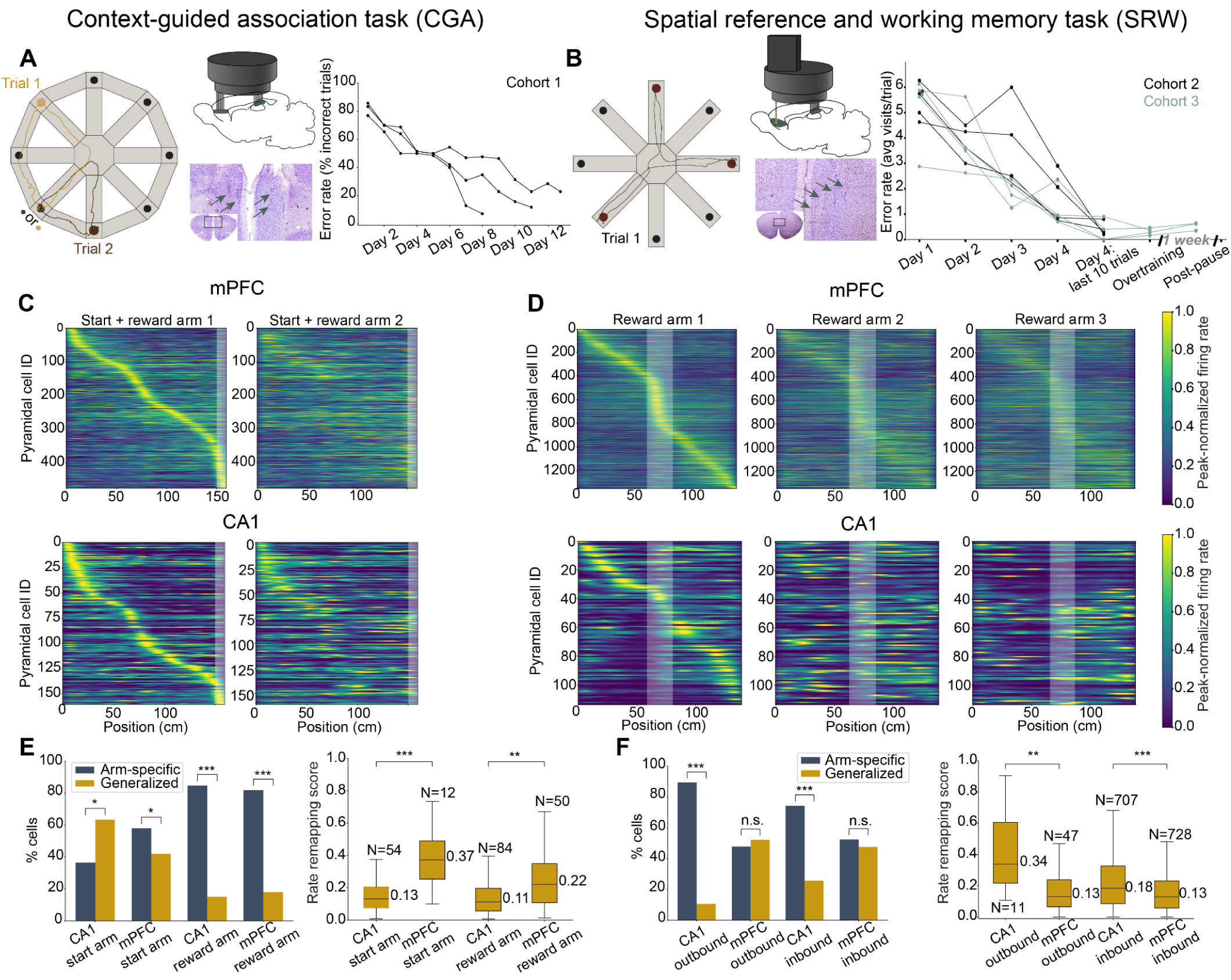
Behavioral paradigms and maze arm-related firing of CA1 and mPFC cells. (**A**) In the CGA task, rats chose reward arms based on food cue identity, with a single cue–arm pairing per trial. *Middle*: Headstage illustration (top) and histology showing tetrode tracks in bilateral mPFC. *Right*: Learning curve. (**B**) In the SRW task, three identical rewards were collected at fixed arms on each trial. *Middle*: Histology shows tracks of the four shanks of a Neuropixels 2.0 probe. *Right*: Learning curve. Cohort 3 continued training on the SRW task and was retested after a pause. (**C–D**) mPFC and CA1 pyramidal neurons show spatial tuning in both tasks. Neurons are sorted by arm 1 peak firing locations. Linearized positions are shown for the entire trial trajectory. Gray shading: reward zones. (**E-F**) Proportion of arm-specific or trajectory-specific (generalized) cells (left), and mean rate remapping score between alternative arms (right) during the CGA task (**E**) and SWR task (**F**). % cells: *** p < 0.001; binomial test; rate remapping: ** p < 0.01, *** p < 0.001; two-sample Kolmogorov-Smirnov test.

We first examined neuronal representations in the session where animals reached optimal performance after learning. Compared to place cells in the hippocampus, mPFC neurons exhibit less spatially restricted activity, frequently generalizing across locations associated with a similar task stage (Hok et al., 2005; Fujisawa et al., 2008; Zielinski et al., 2019; Kaefer et al., 2020). Both regions contained spatially selective cells (Figure 1C, D), consisting of location-dependent, arm-specific neurons and generalized cells with similar firing patterns across multiple arms. The distribution of these coding types differed markedly between the tasks. In the CGA task, a large majority of CA1 and mPFC cells (80%) were arm-specific (Figure 1E left, Figure S2A, C). In the SRW task, the CA1 population consisted primarily of arm-specific cells, whereas the spatially tuned mPFC population was more likely to generalize than in the CGA task, with an even split between arm-specific and generalized coding (Figure 1F left, Figure S2B,D). Context-specific differences were also seen in the start arm during the CGA task, where 40% of CA1 cells and 60% of mPFC cells fired exclusively within a single associational context (Figure 1E left). The remaining generalized cells underwent rate remapping across contexts, which was stronger in mPFC cells than in CA1 cells (Figure 1E right). This rate-remapping trend reversed in the SRW task, favoring CA1 over mPFC cells (Figure 1F right). Together, these findings suggest that mPFC populations adjust to task demands by either specializing or generalizing.

Hippocampal and mPFC spatial representations remap during the acquisition of new spatial tasks (Hollup et al., 2001; Hok et al., 2005, 2007; Dupret et al., 2010), and population activity can drift along a manifold as a session progresses even in the absence of learning (Yang et al., 2024; Kanter et al., 2025). We therefore tested whether this population activity occupied a low-dimensional manifold that simultaneously encoded spatial features (arm identity and position) along with session progression. Using uniform manifold approximation and projection (UMAP) to generate a three-dimensional embedding of SRW task activity, we found that the CA1 manifold clearly differentiated arm identity and spatial position, while capturing session progression to a lesser degree (Figure 2A). By contrast, the mPFC manifold separated session progression and relative arm location, whereas arm-specific representations overlapped within the embedding (Figure 2B). To quantify these relationships, we calculated the structure index (SI) between manifold coordinates and task variables, which quantifies how smoothly task features are topologically organized across the embedding space. Furthermore, by using the projected manifold coordinates, we performed cross-validated decoding of task variables. For the SRW task, these analyses confirmed that the CA1 manifold was organized primarily by arm and position, while session progression was reflected to a lesser extent, though all three variables exceeded shuffled controls (Figure 2C, D). Conversely, mPFC manifolds exhibited a primary organization around relative arm location and session progression rather than specific arm identity. The cross-validated decoding performance remained consistent when expanding the UMAP embedding space to ten dimensions, confirming that the three-dimensional projections sufficiently captured the primary task-relevant variance for these analyses (Figure S3). These results were further validated using a linear embedding of the first three principal components (PCs) from a principal component analysis (PCA), demonstrating that the observed manifold geometries reflected intrinsic population structure rather than non-linear projection distortions. (Figure S4). In the CGA task, similar analyses showed that session progression exhibited the weakest correspondence to the manifold structure (Figure S5, S6), but all three variables could still be decoded significantly better than shuffled controls.

**Figure 2.**
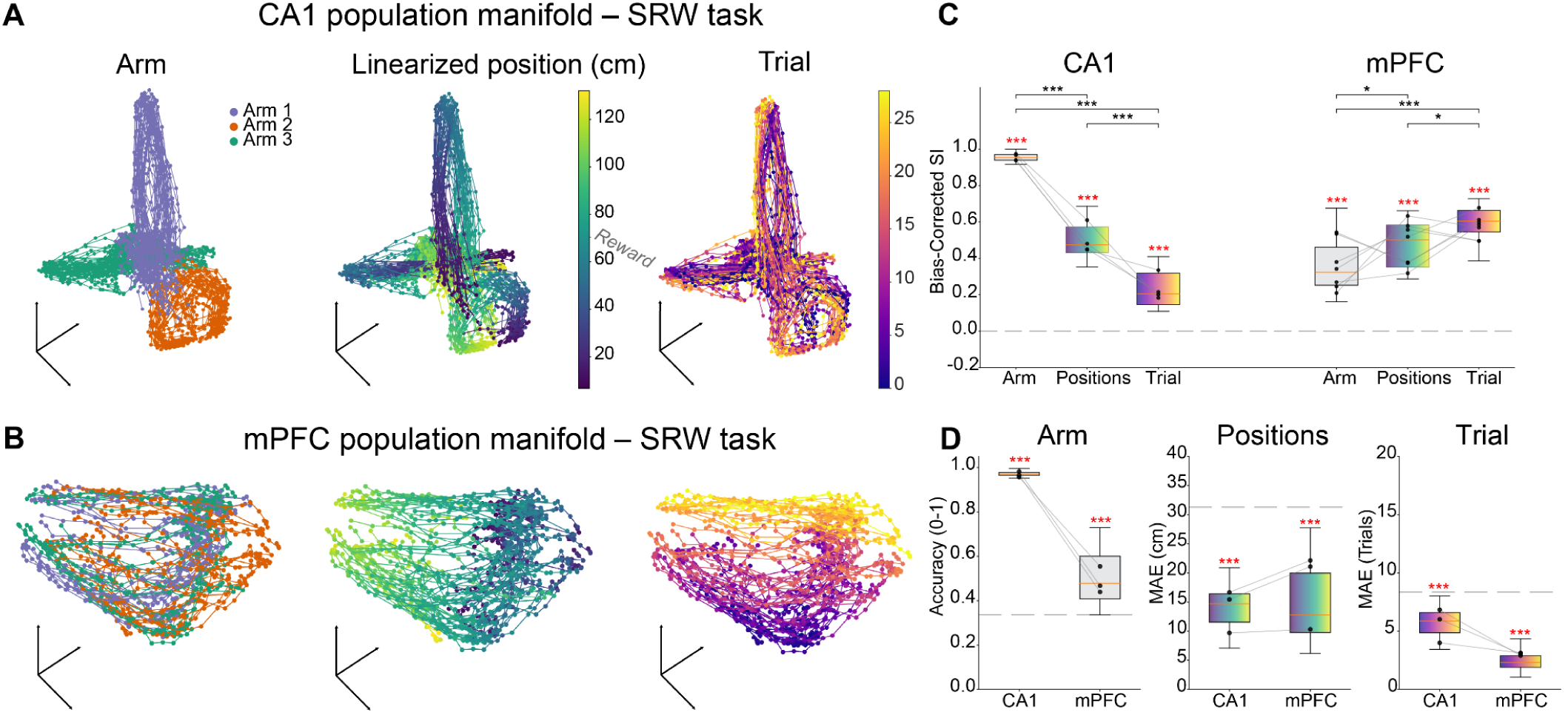
UMAP embedded manifold representation and encoding of task variables in CA1 and mPFC during the SRW task. (**A**) Three-dimensional (3D) UMAP projections of CA1 neural activity for the SRW task colored by (left to right): arm identity, linearized position along the entire trajectory, and trial number. (**B**) 3D UMAP projections of mPFC activity using the same color scheme as in (**A**). (**C**) Bias-corrected structure index (SI) quantifying the spatial organization of task variables within the CA1 and mPFC manifolds. (**D**) UMAP-based decoding performance for arm identity (accuracy), position [mean absolute error (MAE)], and trial progression (MAE). Gray lines: medians for individual animals. Dashed lines: chance levels. Red *: significant differences compared to shuffled distributions, *** *p* < 0.001, Wilcoxon test in (**C**); Mann-Whitney test in (**D**); black *: significant differences between task variables, \**p < 0.05,*\*\*\* *p* < 0.001; Wilcoxon test with Holm-Bonferroni correction.

While these results demonstrated that population manifolds reflected session progression in both regions, aligning with previous observations in CA1 (Yang et al., 2004), how these representations evolved as animals gained task familiarity remained unclear. To address this, we analyzed mPFC data from the second subcohort of animals that continued to perform the SRW task across multiple days after initial learning. As task familiarity increased, mPFC manifolds differentiated arms more effectively, and position decoding was similarly enhanced with experience, while the session progression relationship weakened (Figure 3, Figure S7, S8). A similar reorganization occurred in the CGA task; following the extended training required for task acquisition, the alignment of the UMAP manifolds with session progression decreased (Figure S9). Arm differentiation could not be evaluated in the CGA task because animals frequently visited only one target arm during intermediate learning stages.

**Figure 3.**
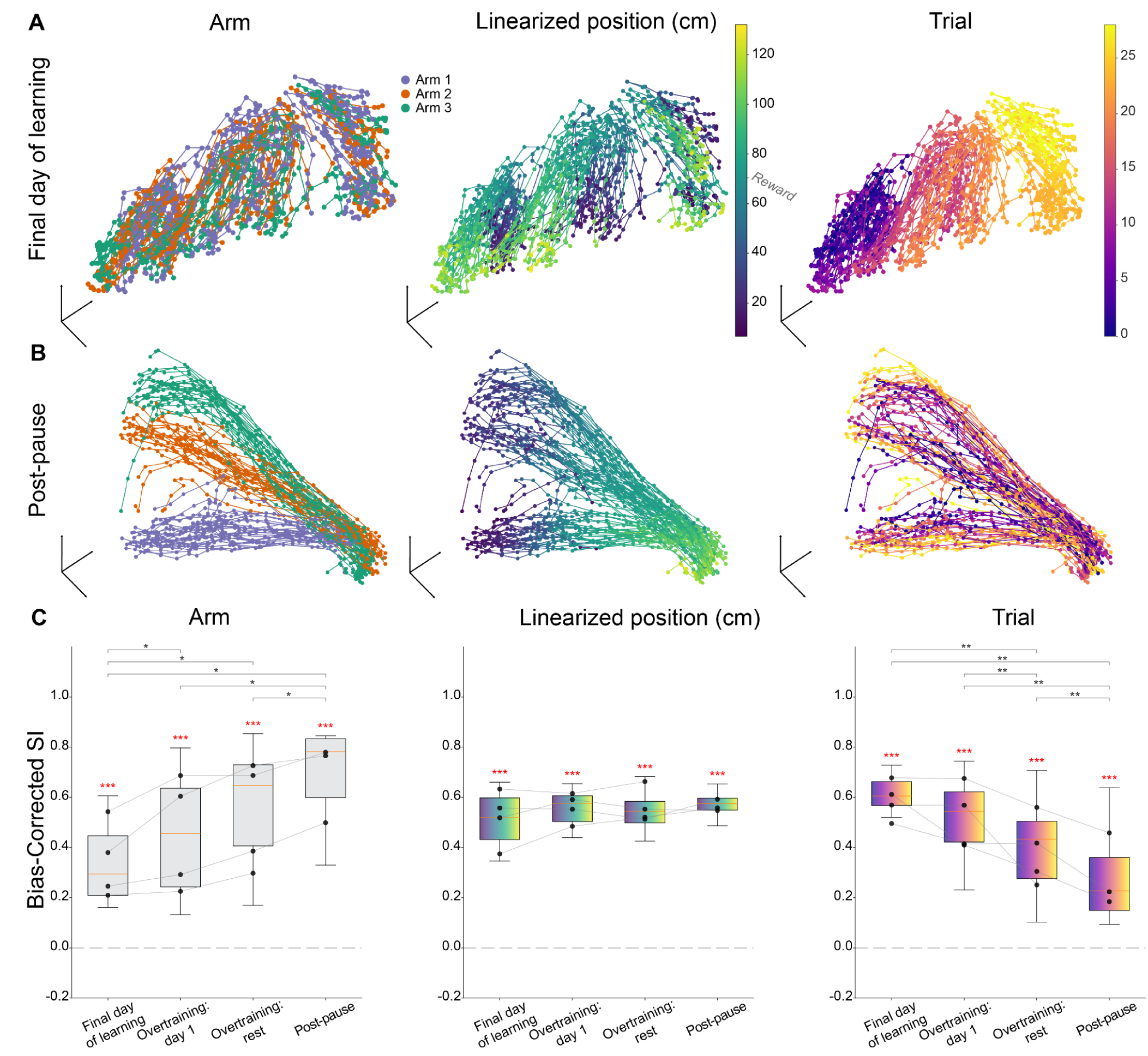
Reorganization of the mPFC manifold with task repetition during the SRW task. (**A-B**) UMAP manifolds during the final learning day (**A**) and post-pause day (**B**). Three-dimensional UMAP projections of mPFC neural activity for the SRW task colored by (left to right): arm identity, linearized position along the entire trajectory, and trial progression. **(C**) Bias-corrected SI tracking the structural organization of task variables across four distinct training phases. Gray lines: medians for individual animals. Dashed lines: chance levels. Red *: significant differences compared to shuffled distributions, *** *p* < 0.001, Wilcoxon test; black *: significant differences between task variables, \**p < 0.05,*\*\* *p* < 0.01; Wilcoxon test with Holm-Bonferroni correction.

Neuropixels recordings enabled longitudinal tracking of individual mPFC neurons to investigate the cellular basis of these representational shifts (Figure 4). We compared coding stability between the final day of learning and a subsequent repeat session, and across a one-week pause in well-trained animals. Under both conditions, approximately 35% of previously non-spatial cells acquired spatial tuning (Figure 4A). These cells initially showed minimal bias in joining the spatial groups, but preferentially joined the arm-specific group in experienced animals (Figure 4B). For neurons that maintained spatial firing across sessions, both arm-specific and generalized cells preferentially retained their respective coding identities. Neurons that preserved their classification across these intervals also exhibited stable firing fields (Figure 4C, Figure S10). A similar trend was seen when the last two days of overtraining were compared before the 1-week pause. (Figure S11). These representational shifts were replicated in terms of the trend of increasing proportion of arm-specific cells when analyzing independent, non-tracked cell populations across each day, confirming that the observed network dynamics were not driven by longitudinal unit-tracking biases (Figure S12). Together, these results indicated that the increased prevalence of arm-specific coding in experienced animals was due to the biased recruitment of previously non-spatial neurons.

**Figure 4.**
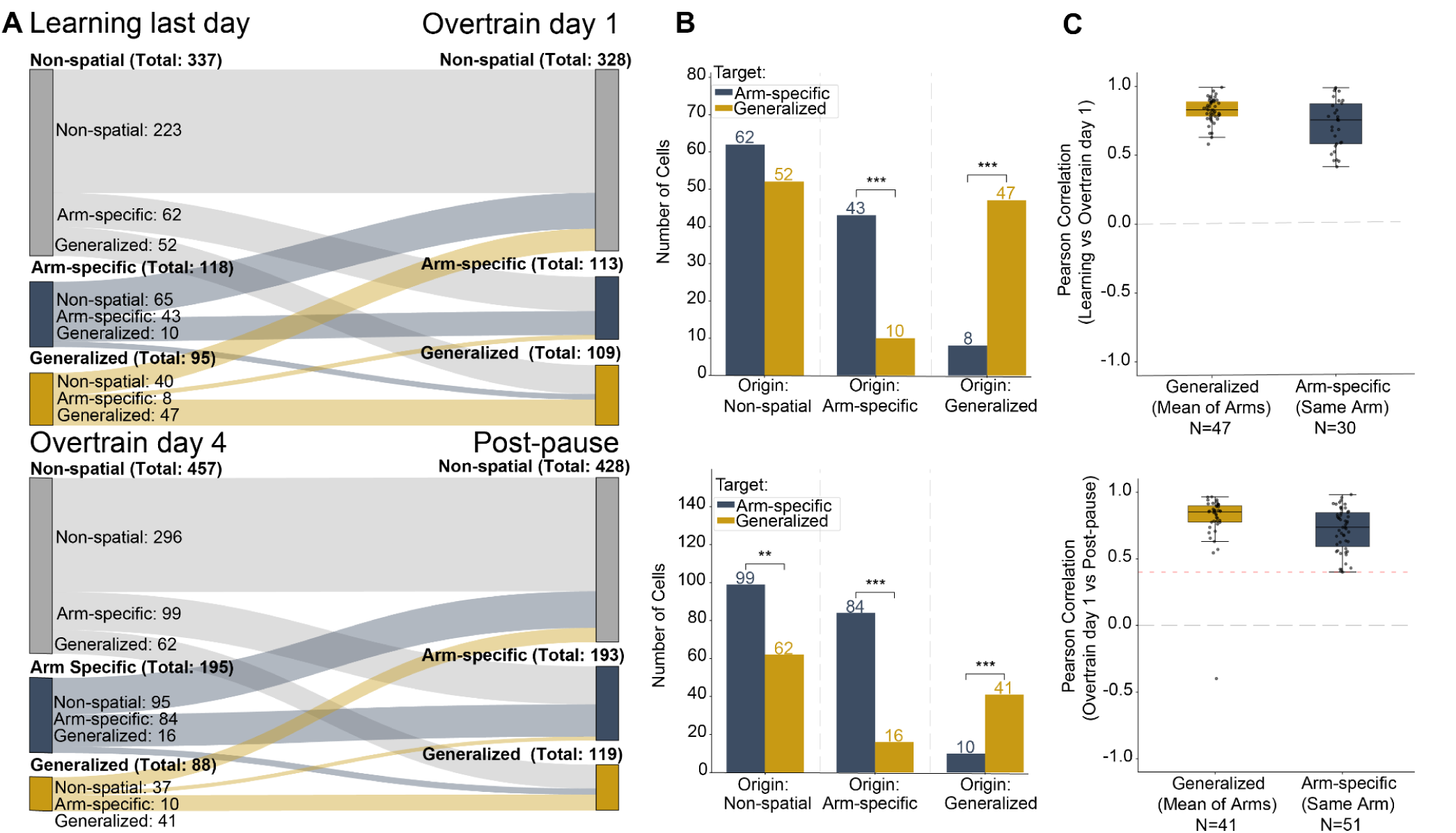
Tracking the coding of individual cells across consecutive recording days. (**A**) Sankey diagrams illustrating the transition of individual mPFC neurons between coding categories (Non-spatial, Arm-Specific, and Generalized) from the final learning day to the first overtraining day (top) and from overtraining day 4 to Post-pause day a week later (bottom). (**B**) Quantification of cell coding changes. Bars indicate the number of cells transitioning into Arm-Specific or Generalized representations, grouped by their original coding state (p < 0.01, *** p < 0.001; binomial test). (**C**) Pearson correlations of spatial rate maps across comparison days for cells that maintained their coding type. Arm-specific neurons are categorized as “Same Arm” if they maintained their coding on the same arm (top: 30 out of the 43 neurons, bottom: 51 out of 84 neurons). The mean correlation across all arms was calculated for the Generalized cells. Black dots: individual cells. For all analyses, each tracked neuron was evaluated independently for outbound (center-to-reward) and inbound (reward-to-center) trajectories.

Across both the SRW and CGA tasks, experienced mPFC manifolds exhibited a decreased correspondence with session progression. However, the improvement in arm differentiation within the SRW task was unexpected, given that prior behavioral and neural data suggested that extended experience instead favors generalized coding (Packard and McGaugh, 1996; Wiltgen and Silva, 2007; Wiltgen and Tanaka, 2013; Morrissey et al., 2017). We hypothesized that this enhanced arm-specific coding reflected the specific requirements for spatial discrimination inherent to the SRW task design. To test this possibility, we analyzed a previously published dataset from a plus-maze task in which animals switched between a spatial strategy targeting a fixed arm and a light-cue-guided strategy (Figure 5, Figure S13). In both conditions, the animals initiated trials from either the north or south start arms and navigated to a fixed or cue-guided destination side arm. Both mPFC and CA1 manifolds differentiated the start arms with similar accuracy across both behavioral strategies (Figure 5D, Figure S14F). However, during the light-cue strategy, side arm differentiation in the mPFC was less accurate than start arm differentiation, whereas CA1 maintained similar accuracy for both arm types (Figure 5C, Figure S14E). These differences in prefrontal representation may reflect the respective task demands, specifically whether tracking specific arm identities was required in executing either of the strategies. Tracking the side arm was not required for successful execution across both strategies, whereas discriminating the start arms remained a requirement during the spatial task.

**Figure 5.**
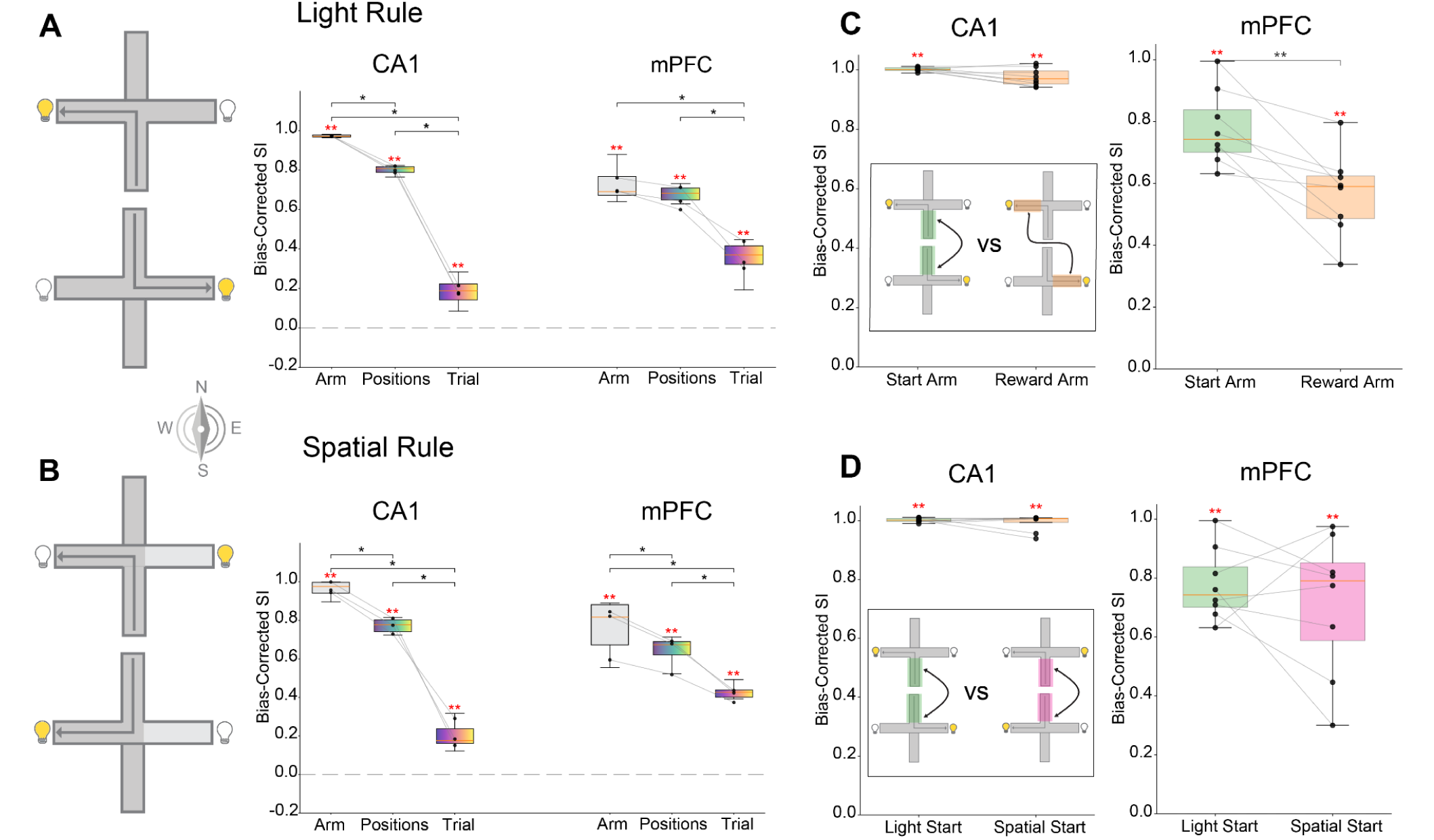
Population manifold encoding across distinct task rules in CA1 and mPFC. (**A-B**) Bias-corrected SI of task variables on UMAP manifolds (see Figure S11) during the Light Rule (**A**) and Spatial Rule (**B**). *Left*: explanation of tasks with the Light and Spatial Rules. *Right*: Bias-corrected SI for arm identity, linearized position, and trial progression for both regions. (**C**) Comparison of SI between the Start Arm and Reward Arm during the Light Rule. (**D**) Comparison of Start Arm SI between the Light Rule and Spatial Rule paradigms. Insets in left panels: illustration of the comparisons. Gray lines: medians for individual animals. Dashed lines: chance levels. Red *: significant differences compared to shuffled distributions ** p < 0.01, Wilcoxon test; black *: significant differences between task variables *p < 0.05,** p < 0.01; Wilcoxon test with Holm-Bonferroni correction.

## Discussion

The primary focus of this work was to investigate the progression of mPFC representations during spatial learning and to examine their refinement as animals gained task proficiency. During the SRW task, we found that projected population activity manifolds initially represented relative locations and advanced with session progression across individual trials, while weakly differentiating individual arms. However, this trend reversed with increasing task familiarity: activity manifolds improved in differentiating arms, whereas their alignment with session progression weakened. Arm differentiation was also observed in the CGA task after a week of training, and on the plus maze, where it varied depending on whether the given arm identity was needed for the performance of tasks executed each day.

Current views of systems consolidation suggest that the mPFC extracts abstract features across repeated experiences, facilitating the formation of generalized knowledge structures such as schemas and gist-like representations(Nadel and Moscovitch, 1997; Frankland and Bontempi, 2005; Tse et al., 2007b, 2011b; McKenzie et al., 2014b; Robin and Moscovitch, 2017; Farzanfar et al., 2023b). Previous behavioral experiments and electrophysiological recordings of mPFC neurons have indeed supported the presence of such abstract coding (Packard and McGaugh, 1996; Yang et al., 2004; Wiltgen and Silva, 2007; Wiltgen and Tanaka, 2013; Yu et al., 2018; Tang et al., 2023; El-Gaby et al., 2024). However, our results suggest that the process is more specialized when animals learn and later practice an entirely novel task over several days, largely because the animal is not simply forming new memories but must establish and refine a new task schema. Our results from the SRW task suggest a two-stage process within the mPFC: we found that initially, a “ready-made,” generalized representation of reward proximity was active, favoring arm-independent trajectory coding. With repeated experience, however, a behaviorally relevant code emerged that differentiated individual maze arms when the task required it. Furthermore, results from the plus-maze strategy-shifting task demonstrated that the mPFC can modulate location representations dynamically in response to task demands, concurrently using both generalized and arm-specific coding across different locations. These findings suggest that cortical schema formation is not a passive averaging process that inevitably discards detail, but rather an optimization process in which the discriminability of representations is shaped by the task at hand (Bernardi et al., 2020).

The finding that session progression was represented to a lesser degree in the population manifolds of experienced animals provides new insights into the mechanism behind learning and consolidation. The mPFC has been suggested to be involved in reinforcement learning-like computations (Ribas-Fernandes et al., 2011; Wang et al., 2018; Baram et al., 2021; Bein and Niv, 2025), as well as adaptive behavior and strategy selection (Milner, 1982; Ragozzino et al., 1999; Floresco et al., 2001; Rich and Shapiro, 2009). Evaluating and updating behavioral choices requires continuously comparing outcomes across consecutive trials. Tracking this trial-by-trial temporal context in early prefrontal representations may be a prerequisite for strategy exploration, enabling the network to maintain the high-dimensional state space necessary for strategy monitoring. Upon task mastery, once a successful behavioral strategy is established, tracking individual trial history becomes computationally redundant. Consistent with the role of the mPFC in reinforcement learning, this process may have indicated an exploration-to-exploitation shift (Kaelbling et al., 1996). Consequently, the network may undergo a form of dimensionality compression by dropping the tracking of session progression in favor of task-relevant variables. Under this framework, the switch from encoding session progression to task variables may signal the acquisition of an effective strategy, indicating that the task has been mastered.

## Acknowledgments

We thank Jago Wallenschus for outstanding technical support. We thank Sofia Taveira and Michele Nardin for comments on the manuscript and Krystsina Masilevich for valuable contributions to figure design. AI tools (Google Gemini Pro 3.1, Claude Opus 4.8) have been used for finding relevant literature and for correcting and editing the text. This work was supported by the Austrian Science Fund (FWF), Cluster of Excellence program “Neuronal Circuits in Health and Disease”.

## Author contributions

Conceptualization: UB, AC, JC

Data collection: UB, AC, KK

Spike sorting and data curation: UB, AC, KK

Data analysis for the paper and figures: UB, AC

Supervision: JC

Writing – original draft: UB, AC, JC

Writing – review & editing: UB, AC, JC

## Data, code, and materials availability

The data will be available from the corresponding author upon request. All custom Python processing scripts along with a representative subset of the dataset are available on GitHub at https://github.com/UladzislauBarayeu/Representational_Dynamics_demo.

## Supplementary Materials

Materials and Methods

## Methods

### Experimental Model and Subject Details

Ten adult male rats (Long Evans, 300-500 g, 3-6 months old) were used in this study. The animals were housed in a separate room under a 12/12 h light/dark cycle and were moved to the recording room during each recording day. They shared a cage with littermates before the surgery. All procedures involving experimental animals were carried out in accordance with Austrian animal law (Austrian federal Law for experiments with live animals) under a project license approved by the Austrian Federal Science Ministry.

### Dataset Overview

- **Dataset 1 (spatial reference and working memory (SRW) task, 8 rats)** 24 CA1 tetrodes + one Neuropixels probe (mPFC)
- **Dataset 2 (cue-guided association (CGA) task, 3 rats)** 32 tetrodes (CA1 + bilateral mPFC)
- **Dataset 3 (plus maze task, 3 rats)** 32 tetrodes (CA1 + bilateral mPFC)

Dataset 3 was previously described in (Kaefer et al., 2020); only key features relevant to the present analyses are summarized below.

### Surgical Procedures

#### Common Surgical Procedures

All animals were anesthetized with isoflurane (0.5–3% in oxygen, 1–2 L/min) and received buprenorphine (0.1 mg/kg) prior to surgery. Six anchoring screws were secured to the skull, with two ground screws positioned over the cerebellum. Following craniotomies and dura removal, tetrode bundles were centered above their respective targets and lowered into the brain. Craniotomies were sealed with paraffin wax, and microdrives were fixed to the skull with dental cement. Postoperative analgesia consisted of meloxicam (5 mg/kg) for up to three days. Animals recovered for one week before tetrodes were gradually advanced in 50–200 µm increments toward their target structures.

### Dataset-Specific Implantation Details

#### Dataset 1 – Tetrodes + Neuropixels

Animals were implanted with a custom microdrive housing:

- 24 individually movable tetrodes targeting bilateral dorsal hippocampal CA1
- A single Neuropixels probe targeting the medial prefrontal cortex (mPFC)

For the first three animals, a Neuropixels 1.0 probe was used; for the 4th animal, a Neuropixels 2.0 probe was used. The Neuropixels probe and the drive with the tetrodes were combined into a single implant in which the probe shank extended ∼3.5 mm deeper than the hippocampal tetrode bundle at implantation. The remaining 4 animals were recorded only from the mPFC with the Neuropixels 2.0 probe.

Tetrodes were constructed from four twisted 12 µm tungsten wires and gold-plated to ∼300 kΩ impedance.

Craniotomies targeted:

- CA1: AP −2.5 to −5.0 mm, ML +-1.5 to +-3.5 mm, DV 1.0 mm
- mPFC: AP +3.0 mm, ML +0.5 mm, DV 4.5 mm

#### Dataset 2 and Dataset 3 – 32 Tetrode Configuration

Animals were implanted with custom microdrives housing 32 individually movable tetrodes arranged in two bundles:

- 16 tetrodes targeting right dorsal CA1
- 16 tetrodes targeting bilateral mPFC (prelimbic area)

mPFC bundles were cut 1–1.5 mm longer than the hippocampal bundle to allow differential depth targeting.

Tetrodes were constructed from four twisted 12 µm tungsten wires and gold-plated to ∼300 kΩ impedance.

Craniotomies targeted:

- CA1: AP −2.5 to −4.5 mm, ML +1.0 to +4.0 mm, DV 1.0 mm
- mPFC: AP 2.5 to 4.6 mm, ML ±0.8 mm, DV 2.3 mm

In these datasets, both hippocampal and prefrontal tetrodes were gradually lowered into their respective target layers after recovery.

### Behavioral Tasks

#### Dataset 1 – SRW Task

Animals were trained on a SRW task in an eight-arm radial maze.

At the start of each trial, rats were confined to the central platform while three fixed arms were baited with 20 mg food pellets. After baiting, all eight doors were opened, and animals were allowed to retrieve all three rewards. The trial ended once all rewards were collected.

Daily sessions consisted of eight trials separated by 2-minute inter-trial intervals. Sleep was recorded immediately before and after each behavioral session.

On day four, training continued until animals reached the learning criterion (≥80% correct arm visits).

For the four animals, overtraining recordings were performed, consisting of 30 trials per session, with two sessions per day for three consecutive days. After this period, recordings were paused for one day and then resumed for an additional two sessions under the same task configuration. Subsequently, a one-week break was introduced, after which recording resumed under the same conditions for one more day.

#### Dataset 2 – CGA Task

Following recovery and habituation, training began once animals were able to complete habituation trials within 2–5 minutes, and tetrodes had reached their target locations. Each training day consisted of alternating sleep and behavioral blocks: 40–60 minutes of sleep, followed by a training block, another 40–60 minutes of sleep, a second training block, and a final 40–60 minutes sleep session.

In the cue-guided association (CGA) task, rats were given a food cue in a designated start arm of an eight-arm maze. The identity of the food cue indicated which of the remaining arms was rewarded, with two paired cue-arm associations trained per animal, and the rewarded arms remaining constant across training.

During each trial, the rat began in the start arm and received a small food cue placed in a reward well at the end of the arm (either 1/4 chocolate chip, 1/4 Kellogg’s Honey Loop, or one sunflower seed). After consuming the cue, access to the center of the maze was opened while the outer ring remained blocked. Upon entering the center and selecting one of the goal arms, the door behind the animal was closed. The rat was permitted to poke its head into the chosen arm; once the whole body entered, the door was fully closed.

Only one arm contained a reward. If the animal selected the correct arm, it received a larger reward of the same type as the cue (1 chocolate chip, 1 Honey Loop, or 3 sunflower seeds; reward weights were approximately matched). Additionally, a sucrose pellet was placed in the start arm on correct trials to incentivize return to the start location.

Two cue types were presented in parallel during training, and their order was randomized. Rewarded arms and initial cue types (two out of three possible cues) were counterbalanced across animals. The cue–arm associations remained constant throughout the task.

On the first training day, training targeted 30 trials, with animals completing 18–30 trials. On subsequent days, animals completed 26–60 trials, and from day four onward, all animals performed 70 trials per day. Training sessions were split into two halves separated by a sleep period. Sessions typically lasted 1–3 hours, though occasionally extended longer (up to ∼5 hours); sessions were terminated early if animals showed signs of fatigue or reduced motivation.

Once animals reached a performance criterion of ≥80% correct trials, an additional stabilization day was performed.

#### Dataset 3 – Plus Maze Task (Previously Published)

Dataset 3 employed a rule-switching on an elevated plus maze; see (Kaefer et al., 2020) for full details.

Briefly, animals initiated trials from one of two start arms and were required to retrieve reward according to either:

- A spatial rule (reward always in a fixed goal arm), or
- A response/light rule (reward in the illuminated arm).

Rule changes were not signaled and required trial-and-error learning. Behavioral blocks consisted of pre-switch, switching, and post-switch phases. For UMAP analysis, only data from pre-switch and post-switch blocks with the light task were analyzed.

Performance criteria were adjusted for spatial and light rules to equate trajectory sampling.

### Histology and Reconstruction of Recording Positions

At the end of experiments, animals were deeply anesthetized with an overdose of pentobarbital (300 mg/ml) and transcardially perfused with 0.9% saline followed by 4% formaldehyde. For CGA animals, tetrode locations were marked by injecting a continuous, bipolar current at 100 µA for 30 seconds immediately before the perfusion. Brains were extracted and post-fixed in 4% formaldehyde, then cryoprotected in 30% sucrose at 4 °C until they sank. Tissue was rapidly frozen and sectioned coronally on a cryostat (50–60 µm), mounted on glass slides, and Nissl-stained (cresyl violet). Neuropixels probe and tetrode tracks were identified from the stained sections. For hippocampal tetrode recordings, datasets were included only when sharp-wave ripples (SWRs) were evident in the local field potential.

### Data Acquisition

Across all datasets, extracellular signals were recorded during task performance and sleep sessions. Animal position was tracked using head-mounted LED clusters monitored by an overhead camera. Prior to each recording day, CA1 and mPFC tetrodes were adjusted to optimize single-unit yield.

#### Dataset 1 – Neuropixels + Tetrode Configuration

Neural activity was recorded simultaneously from tetrodes in CA1 and a Neuropixels probe in medial prefrontal cortex (mPFC).

For Neuropixels 1.0 probes (Jun et al., 2017), the 384 electrodes closest to the probe tip were enabled. For Neuropixels 2.0 probes (Steinmetz et al., 2021), a subset of dorsal recording sites was selected to target neurons in prelimbic (PL) and anterior cingulate cortex (ACC). Tetrode signals were amplified using an Intan RHD2132 headstage. All data were acquired with the Open Ephys GUI (Siegle et al., 2017). Neuropixels, tetrode, and video streams were synchronized via TTL pulses generated by the behavioral camera.

Animal position was tracked using a head-mounted RGB LED cluster (red, green, blue).

#### Dataset 2 and Dataset 3 – 32 Tetrode Configuration

Extracellular signals from tetrodes were pre-amplified using a headstage (4 × 32 channels, Axona Ltd., UK). Local field potentials and multi-unit activity were continuously digitized at 24 kHz using a 128-channel Axona data acquisition system.

Two differently sized infrared LEDs mounted on the preamplifier headstage were used for positional tracking. Before each recording session, hippocampal tetrodes were advanced to optimize unit yield from the CA1 pyramidal layer. In addition, mPFC tetrodes were lowered daily by approximately 30–50 µm to record from a new population of neurons across sessions.

### Spike detection and sorting

For hippocampal tetrode data, spike detection and automated sorting were performed with MountainSort (Chung et al., 2017), whereas Neuropixels data were sorted with Kilosort 4.0 (Pachitariu et al., 2024). All putative units underwent manual curation in Phy2(https://github.com/cortex-lab/phy), and were classified as putative pyramidal cells or interneurons based on spike waveform features, autocorrelograms, and mean firing rates (Csicsvari et al., 1999).

### Longitudinal Unit Tracking Across Sessions

To track individual mPFC neurons across multi-day intervals, binary streams from paired sessions were independently drift-corrected and temporally concatenated. Spike sorting was performed on the composite recording using Kilosort 4.0 with automated drift correction disabled to preserve pre-aligned spatial baselines. Tracked units were manually curated in Phy2 to confirm stable waveform geometries, consistent channel footprints, and uncontaminated composite autocorrelograms across the session junction.

### Linearization of Position

Two-dimensional position (video tracking) was linearized separately for each arm of the radial maze by projecting each sample onto the arm’s centerline and converting to arc-length (distance along the centerline). Only baited arms were included in analyses. For the CGA task, the linearized trajectory spanned from the start arm through the central segment into the baited arm, yielding a total path length of 160 cm (70 cm start arm, 20 cm center platform, 70 cm baited arm). For the SRW task, three baited arms were analyzed, with outbound and inbound paths combined onto a single linear axis of 140 cm and the reward located at the midpoint of the axis.

### Rate map construction

Analyses were restricted to putative pyramidal cells that fired ≥1 spike in ≥60% of trials. For each trial, the linearized position was binned at 1-cm resolution. In each bin, spikes were counted and divided by the animal’s occupancy time to obtain the firing rate. Only time points with running speed > 5 cm/s contributed to occupancy and spike counts. Raw rate maps were smoothed with a Gaussian kernel (σ = 3 cm). Spatial tuning was quantified using Skaggs’ spatial information (Skaggs et al., 1993).

### Spatial Tuning and Classification of Task-Related Neurons

To characterize the representation of spatial trajectories in the CA1 and mPFC, we analyzed neural activity during two distinct task phases: *Going to Reward* (linearized bins 0–70) and “Return to Center” (bins 70–end) for the SRW dataset. For the CGA dataset, “Common arm” (linearized bins 0–70) and “Reward arm” (bins 70–end). Cells were first filtered based on their spatial information content. A neuron was included in further analysis if its spatial information exceeded a threshold of 0.1 bits/spike on at least one of the three arms. Furthermore, neurons were only analyzed for a specific task phase if their peak firing rate occurred within the spatial boundaries of that window.

### Definition of Arm-Specific and Generalized Representations

We classified spatially tuned neurons into two functional categories—Arm-specific or Generalized—based on the similarity of their spatial firing profiles across the three arms. For each cell, we calculated the pairwise Pearson correlation coefficients (r) between the firing rate vectors of the three arms within the relevant task window.

- Generalized Cells: Neurons were classified as generalized if at least two pairwise correlations between arms exceeded a threshold of r = 0.6. These cells maintain a similar spatial representation across different trajectories.
- Arm-specific Cells: Neurons were classified as arm-specific if the maximum pairwise correlation across arms was below the r = 0.6 threshold. In longitudinal analyses, tracked arm-specific cells were classified as retaining their physical arm preference (*Same Arm*) if their cross-session rate map correlation exceeded r=0.4.
- Rest: Neurons failing to meet the spatial information threshold or the peak activity location criteria were excluded from the specific representation analysis.

### Quantification of Rate Remapping

To investigate magnitude-based changes in firing (rate remapping) within stable spatial representations, we calculated a Rate Remapping Score for all qualifying pairs of arms in Generalized cells (where pairwise r > 0.6). The score was defined as:

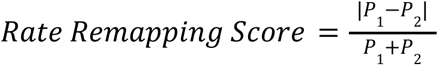

where *P_1_* and *P_2_* represent the peak firing rates on the two compared arms. This index ranges from 0 (no remapping) to 1 (complete rate remapping).(Leutgeb et al., 2005)

### Low-Dimensional Embedding and Decoding Analysis

Neural spiking activity recorded during maze behavior was converted to spike counts using non-overlapping 300-ms time bins. To isolate task-relevant activity, data were filtered by a predefined minimum running speed threshold. For each session, this produced a population activity matrix

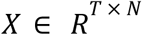

where *T* denotes the number of time bins and *N* the number of recorded neurons; each row corresponds to the population firing vector at a single time bin. To reduce high-frequency noise while preserving local temporal structure, spike counts were smoothed using a 5-bin moving average. Crucially, this smoothing was applied strictly within contiguous behavioral trial segments to prevent smoothing across distinct spatial events or trial boundaries.

Behavioral labels (arm identity, spatial position, and trial number) were aligned to each time bin.

To perform statistical comparison across learning stages, each recording session during the SRW and CGA tasks was divided into two halves (early and late) based on the total number of behavioral trials. Population activity from these early and late subgroups was embedded and analyzed independently. This was performed for the Structure Index and cross-validated decoding comparisons.

#### UMAP and PCA Dimensionality Reduction

Population activity was embedded into a low-dimensional manifold using UMAP (McInnes et al., 2020). UMAP was applied with the following parameters: n_neighbors = 50, min_dist = 0.1, n_components = 3, metric = ’euclidean’.

The algorithm constructs a weighted k-nearest-neighbor graph in the high-dimensional firing-rate space and optimizes a low-dimensional representation that preserves local neighborhood structure while maintaining global manifold organization.

For linear comparisons, PCA was also applied to project the same smoothed population activity into 3 principal components.

Each point in the resulting 3-dimensional embedding corresponded to the neural population state during a single 300-ms time bin.

#### Structure Index of the Manifold

To quantify how smoothly task variables were organized across the neural manifold, we computed the Structure Index (SI) (Sebastian et al., 2024). This metric was evaluated on the manifolds and rigorously bias-corrected using a shuffle-label null distribution.

For continuous variables (position and trial progression), we computed the Neighborhood Time Predictability Index. For each point in the embedding, its k-nearest neighbors (k=20) were identified in the low-dimensional space. The predicted value *t̂* was calculated as the mean label of these neighbors, and the SI was calculated as:

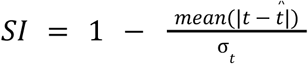

where *t* is the true continuous label, and σ_t_ is the standard deviation of the label distribution.

For categorical variables (arm identity), we computed the Neighborhood Category Predictability Index. The local agreement was defined as the proportion of the 20 nearest neighbors sharing the same categorical label as the center point. The SI was calculated as:

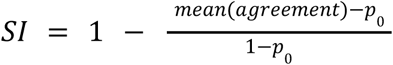

where *p*_0_ represents the chance-level agreement based on the squared class probabilities.

To account for inherent data structure and finite sample biases, all SI values were bias-corrected by subtracting the mean metric obtained from 50 permutations of shuffled behavioral labels.

#### Cross-Validation Decoding from the Low-Dimensional Manifold

To evaluate the explicit decodability of task variables, decoding analyses were performed using the 3- or 10-dimensional embedding coordinates as features. To ensure robust evaluation and prevent temporal data leakage, all decoding analyses utilized a 5-fold Stratified Group

Cross-Validation procedure. The data was split such that all time bins belonging to a single continuous event (e.g., one complete pass through an arm) were kept together in either the training or the test set.

For each fold, the manifold mapping (UMAP or PCA) was fit exclusively on the training data (80%), and the test data (20%) was projected into this newly learned embedding space:

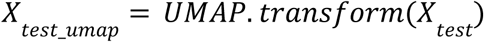

This procedure ensured that the test data did not influence manifold construction, preserving a proper assessment of generalization.

All task variables were decoded using k-nearest-neighbor (KNN; k=5) models trained on the low-dimensional coordinates of the training set and evaluated on the held-out test set. Arm identity was decoded using a KNN Classifier(https://scikitlearn.org/stable/modules/generated/sklearn.neighbors.KNeighborsClassifi <u>er.html</u>), with performance quantified as mean classification accuracy. Continuous variables (linearized position and trial index) were decoded using a KNN

Regressor(https://scikit-learn.org/stable/modules/generated/sklearn.neighbors.KNeighborsRegressor.html), because the temporal gradient is often nested within the spatial manifold topology, making local topological decoders highly effective. Continuous decoding performance was quantified using Mean Absolute Error (MAE). All decoding performances were compared against chance levels derived from identical models trained on shuffled target labels.

### Software and Code Declarations

All custom neural data processing, dimensionality reduction pipelines, and cross-validated decoding workflows were executed in Python. Large language models (specifically Gemini Pro 3.1, Google) were utilized for code drafting, syntax adaptation, and debugging assistance. All AI-assisted code was manually inspected, tested, and validated by the authors, who assume absolute responsibility for the accuracy, reproducibility, and integrity of the analytical codebase.

### Supplementary figures

**Supplementary Figure S1.**
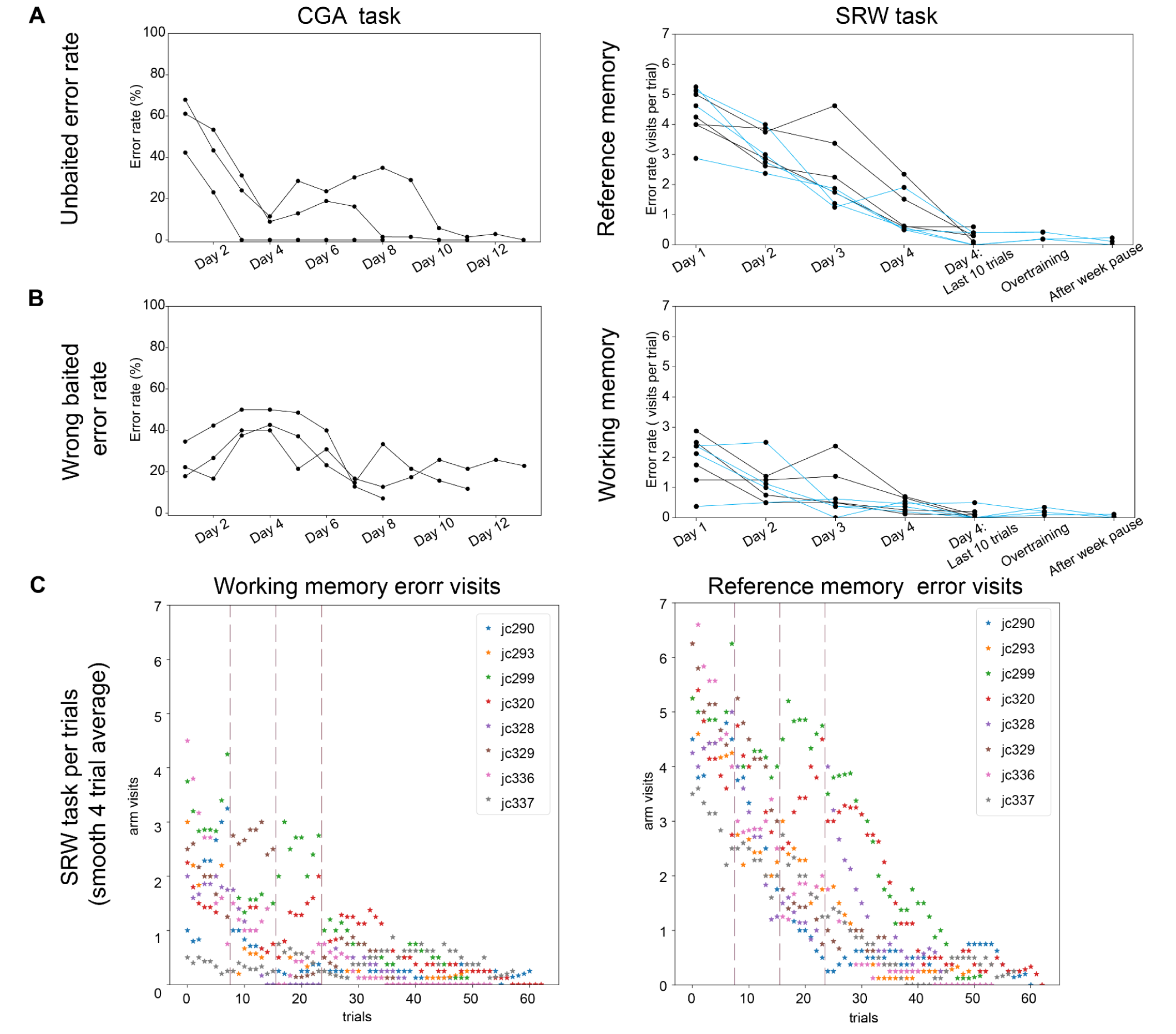
Trial-by-trial error rates. (**A, B)** *Left:* Errors in the CGA task. Performance across 13 training days (n = 3). *Right:* Errors in the SRW task. (**A**) *Left:* Unbaited error rate represents visits to arms that are never rewarded across any context. *Right:* reference memory error (visit an arm without reward) (**B)** *Left:* Wrong baited error rate (%) represents visits to rewarded arms that do not match the specific food cue of the current session. *Right:* Working memory error (%) represents visits to the same arms more than once. (**C**) Working and reference memory errors per trial for individual animals during SRW task, smoothed with a moving average over four trials. Dashed lines: training day boundaries. Individual subjects are color-coded to illustrate varying rates of task acquisition and error reduction.

**Supplementary Figure S2.**
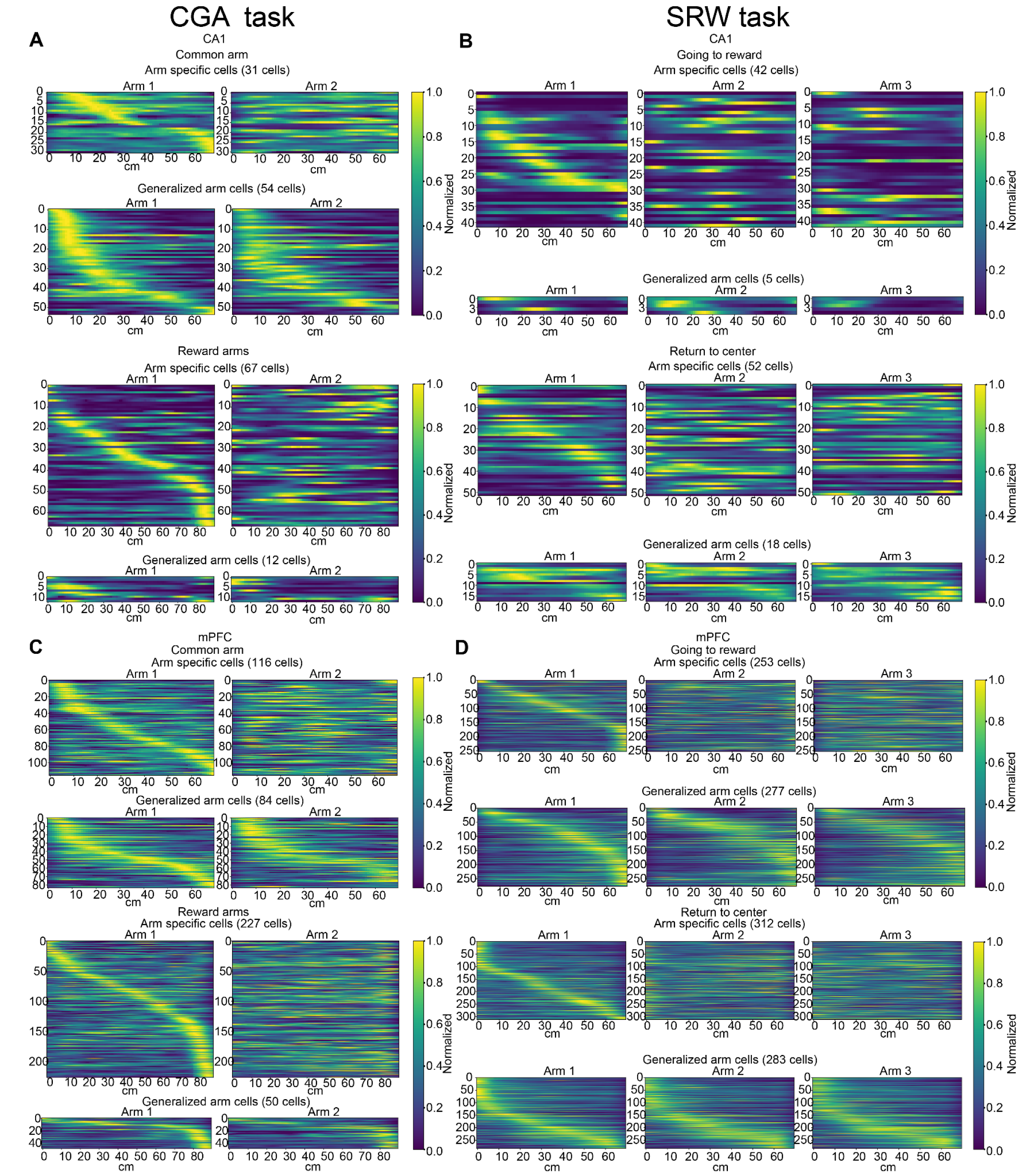
Spatial rate maps for CA1 and mPFC. (**A**, **C**) CGA task. Normalized firing rates for CA1 (**A**) and mPFC (**C**) neurons. Cells are categorized by activity in the common start and reward arms. Neurons are classified as arm-specific or generalized based on their spatial selectivity across contexts. Common arm rate maps were differentiated based on whether the animal would go to arm1 or arm2 next. (**B**, **D**) SRW task. Firing rates for CA1 (**B**) and mPFC (**D**) during outbound (center to reward) and inbound (return to center) trajectories.

**Supplementary Figure S3.**
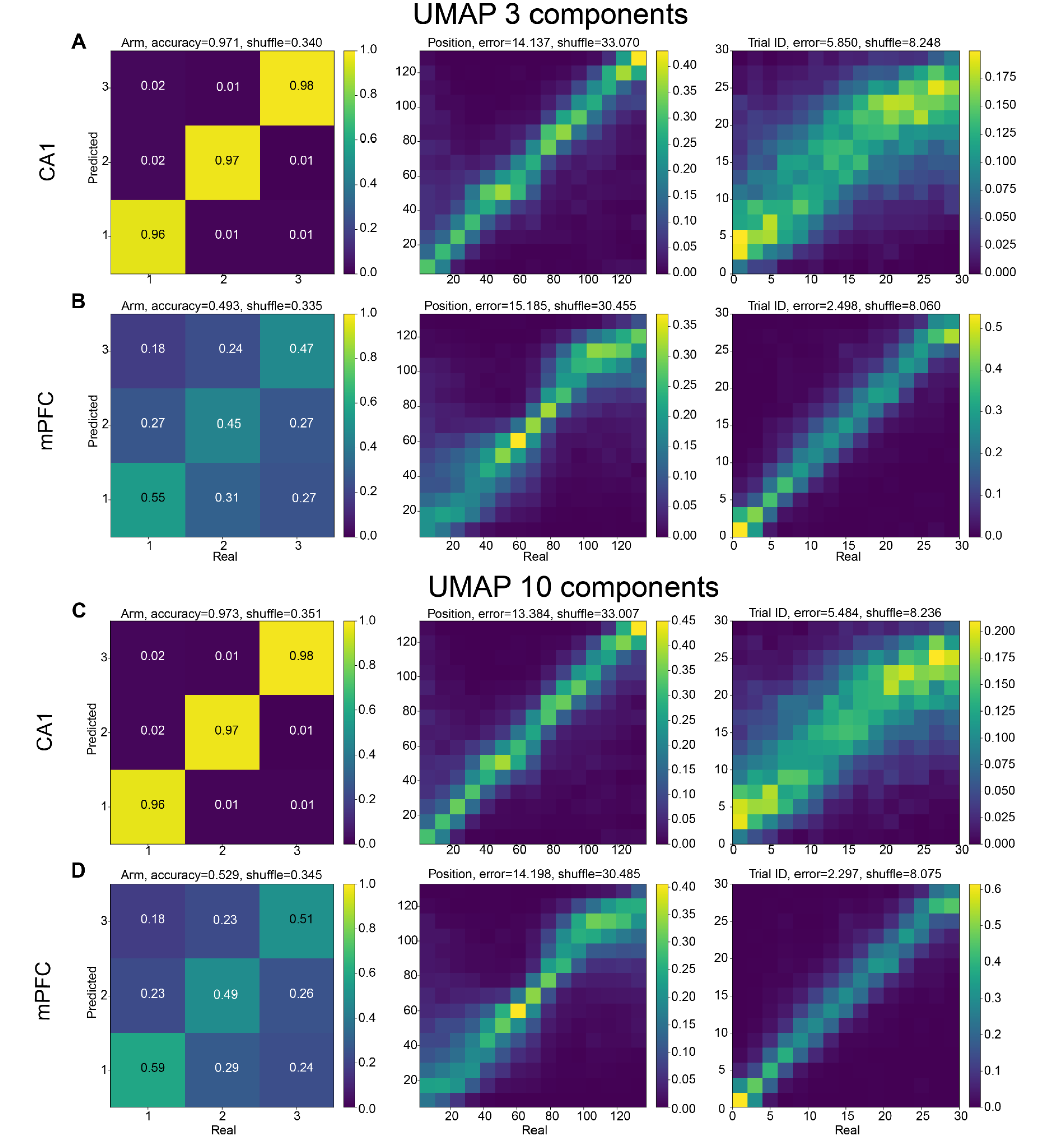
Cross-validated decoding of task variables from three and ten-dimensional UMAP manifolds. (**A**, **B**) Decoding results from UMAP projections of CA1 (**A**) and mPFC (**B**) population activity during the SRW task. Left to right: confusion matrices for arm identity, and joint distributions of predicted versus real linearized position and trial progression. (**C**, **D**) Corresponding decoding profiles from 10-dimensional UMAP projections for CA1 (**C**) and mPFC (**D**) populations. Values above plots: overall model accuracy (arm identity) or mean absolute error (position and trial progression), alongside chance levels from shuffled distributions. Data were pooled together across all animals. Detailed performance distributions correspond to summaries in Figure 2D.

**Supplementary Figure S4.**
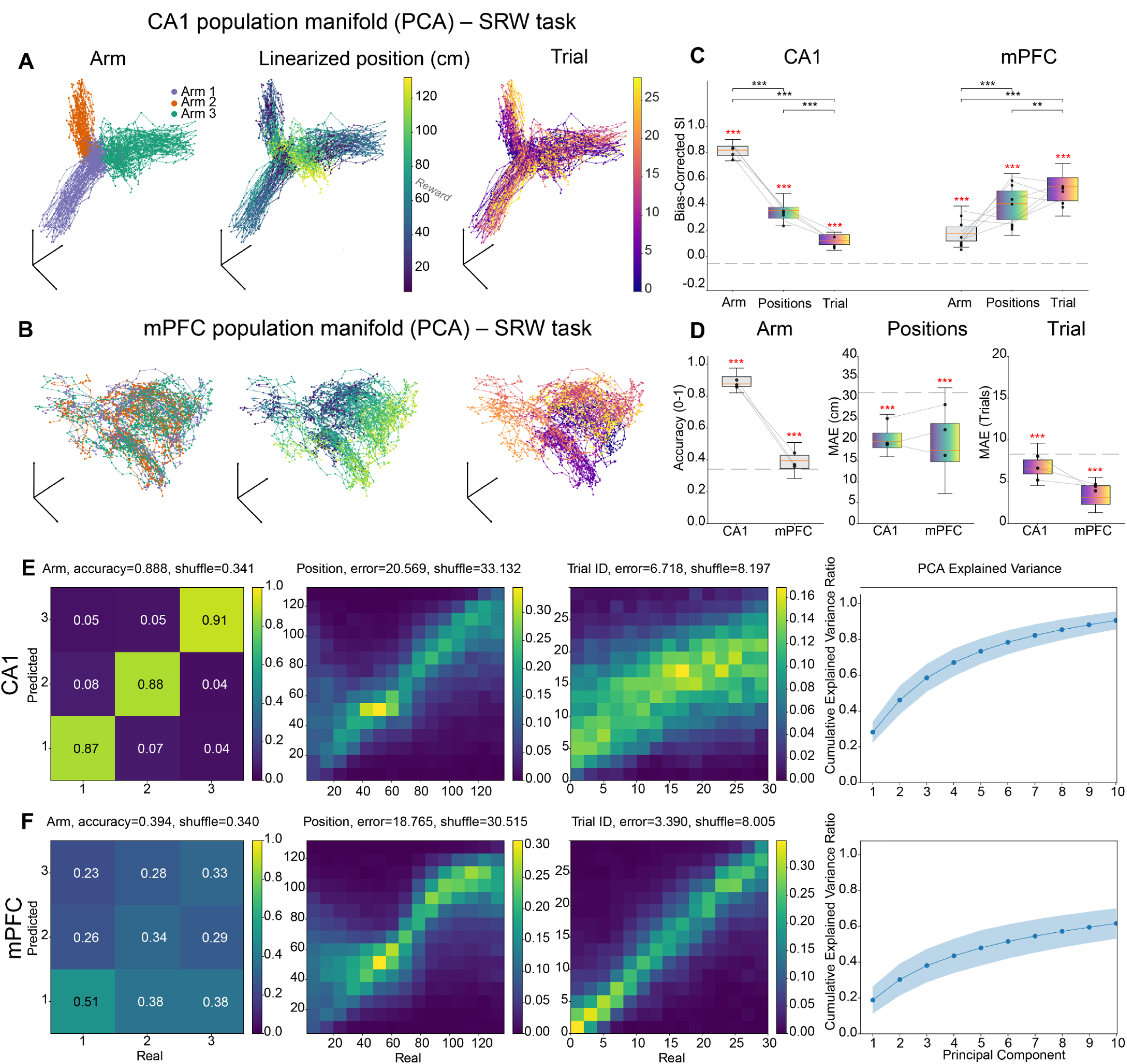
PCA-derived manifold representation and encoding of task variables during the SRW task. (**A**) 3D PCA projections of CA1 neural activity for the SRW task, colored by (left to right): arm identity, linearized position along the entire trajectory, and trial number. (**B**) 3D PCA projections of mPFC activity using the same color scheme as in (**A**). (**C**) Bias-corrected SI quantifying the spatial organization of task variables within the CA1 and mPFC manifolds. (**D**) PCA-based decoding performance for arm identity (accuracy), position (MAE), and trial progression (MAE). (**E**, **F**) Left columns: Decoding results from PCA projections of CA1 (**A**) and mPFC (**B**) population activity during the SRW task. Left to right: confusion matrices for arm identity, and joint distributions of predicted versus real linearized position and trial progression. Data were pooled together across all animals. *Right panels:* cumulative fraction of explained variance across the first 10 principal components (mean ± SEM across sessions). For (**C**) and (**D**), significance markers and statistical tests are identical to those described in Figure 2.

**Supplementary Figure S5.**
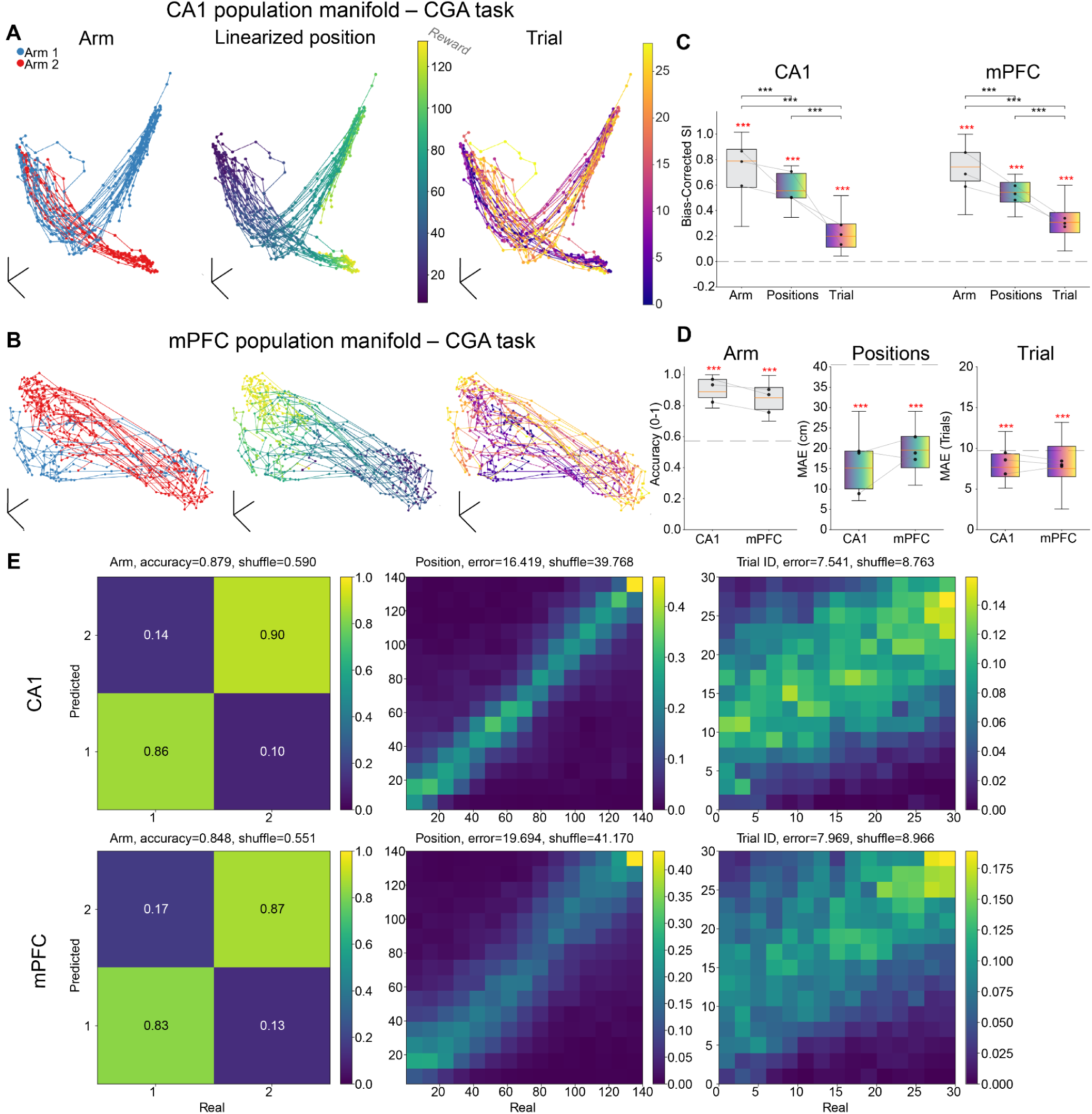
UMAP-manifold representation and decoding of task variables during the CGA task. (**A**) 3D UMAP projections of CA1 neural activity for the CGA task colored by (left to right): arm identity, linearized position along the entire trajectory, and trial number. (**B**) 3D UMAP projections of mPFC activity using the same color scheme as in (**A**). (**C**) Bias-corrected SI quantifying the spatial organization of task variables within the CA1 and mPFC manifolds. (**D**) UMAP-based decoding performance for arm identity (accuracy), position (MAE), and trial progression (MAE). (**E**) Decoding results from UMAP projections of CA1 (**A**) and mPFC (**B**) population activity during the CGA task. Left to right: confusion matrices for arm identity, and joint distributions of predicted versus real linearized position and trial progression. For confusion matrices, data were pooled together across all animals. For (**C**) and (**D**), significance markers, and statistical tests are identical to those described in Figure 2.

**Supplementary Figure S6.**
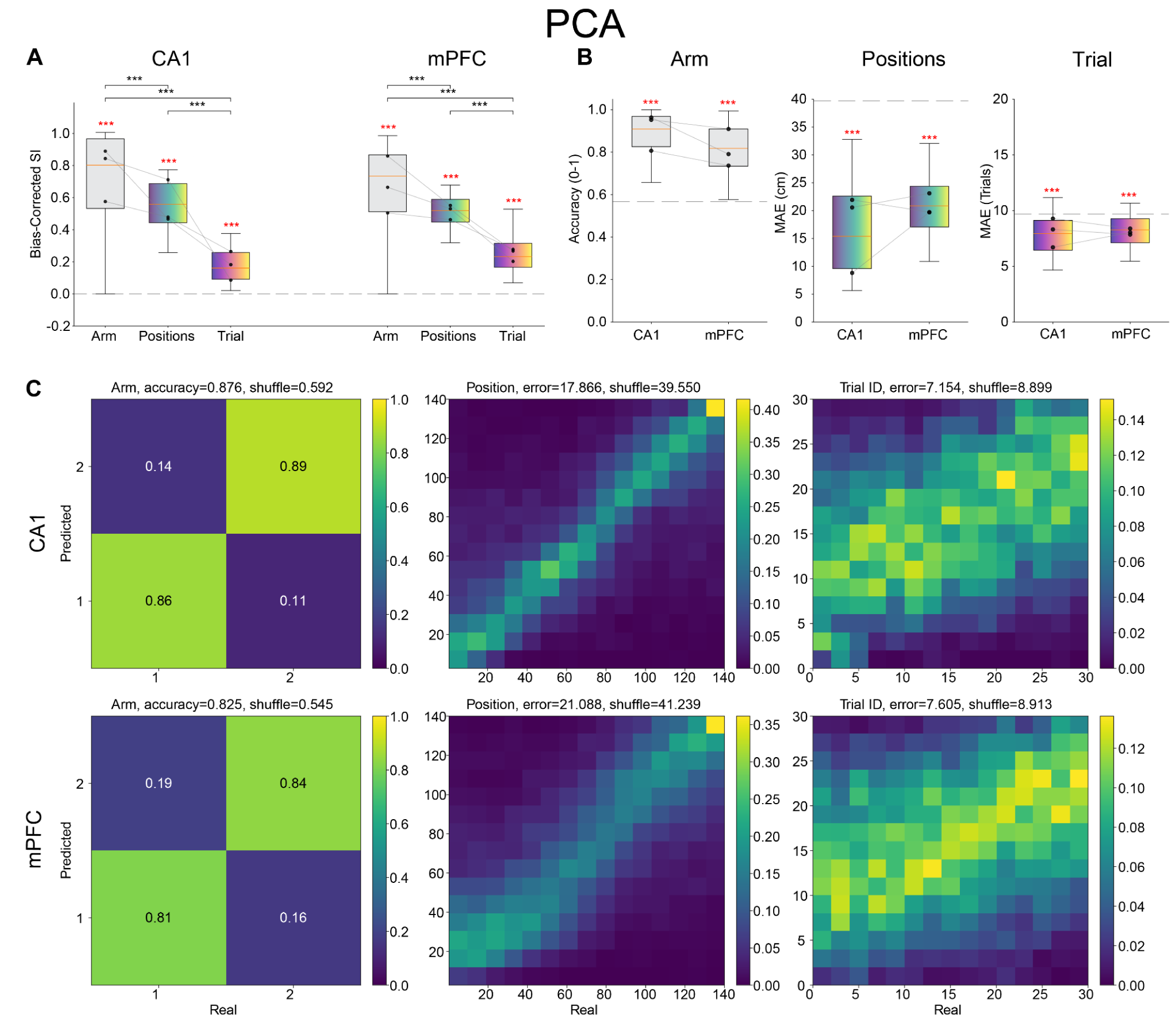
PCA-derived manifolds and decoding of task variables during the CGA task. (**A**) PCA-derived bias-corrected SI quantifying the spatial organization of task variables within the CA1 and mPFC manifolds. (**B**) PCA-based decoding performance for arm identity (accuracy), position (MAE), and trial progression (MAE). (**C**) Decoding results from PCA projections of CA1 and mPFC population activity during the CGA task. Left to right: confusion matrices for arm identity, and joint distributions of predicted versus real linearized position and trial progression. For confusion matrices, data were pooled together across all animals. For (**A**) and (**B**), significance markers and statistical tests are identical to those described in Figure 2.

**Supplementary Figure S7.**
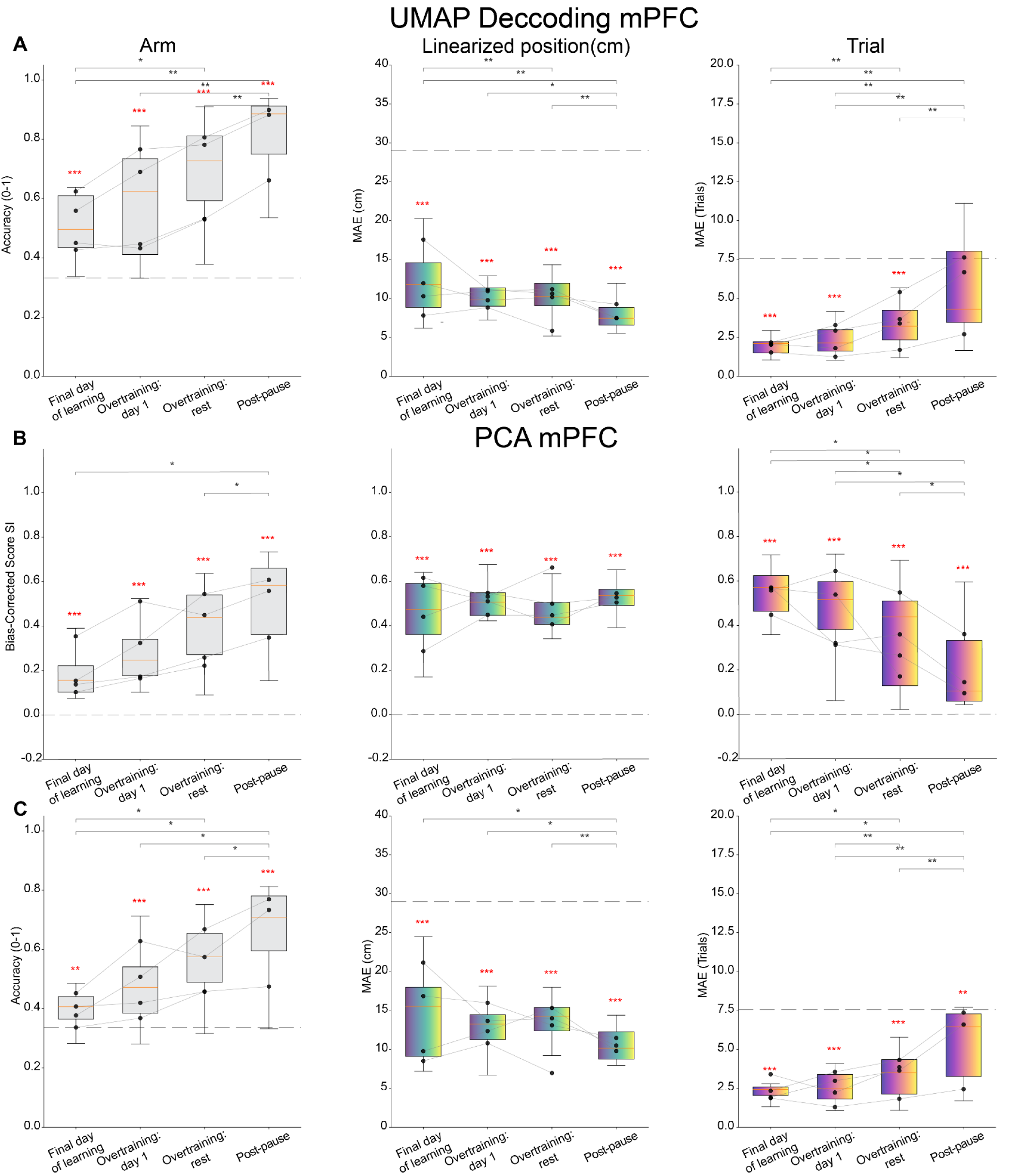
Manifold analysis and decoding of task variables with task repetition during the SRW task. (**A**) mPFC UMAP decoding for arm identity (accuracy), linearized position (MAE), and trial progression (MAE) across training phases (Fig. 3). (**B**) Bias-corrected SI for these variables from linear PCA projections. (**C**) PCA-based decoding for the same variables. Gray lines: medians for individual animals. Dashed lines: chance levels. Red *: significant differences compared to shuffled distributions, *** p < 0.001, Wilcoxon test; black *: significant differences between task variables, *p < 0.05,** p < 0.01; Wilcoxon test with Holm-Bonferroni correction.

**Supplementary Figure S8.**
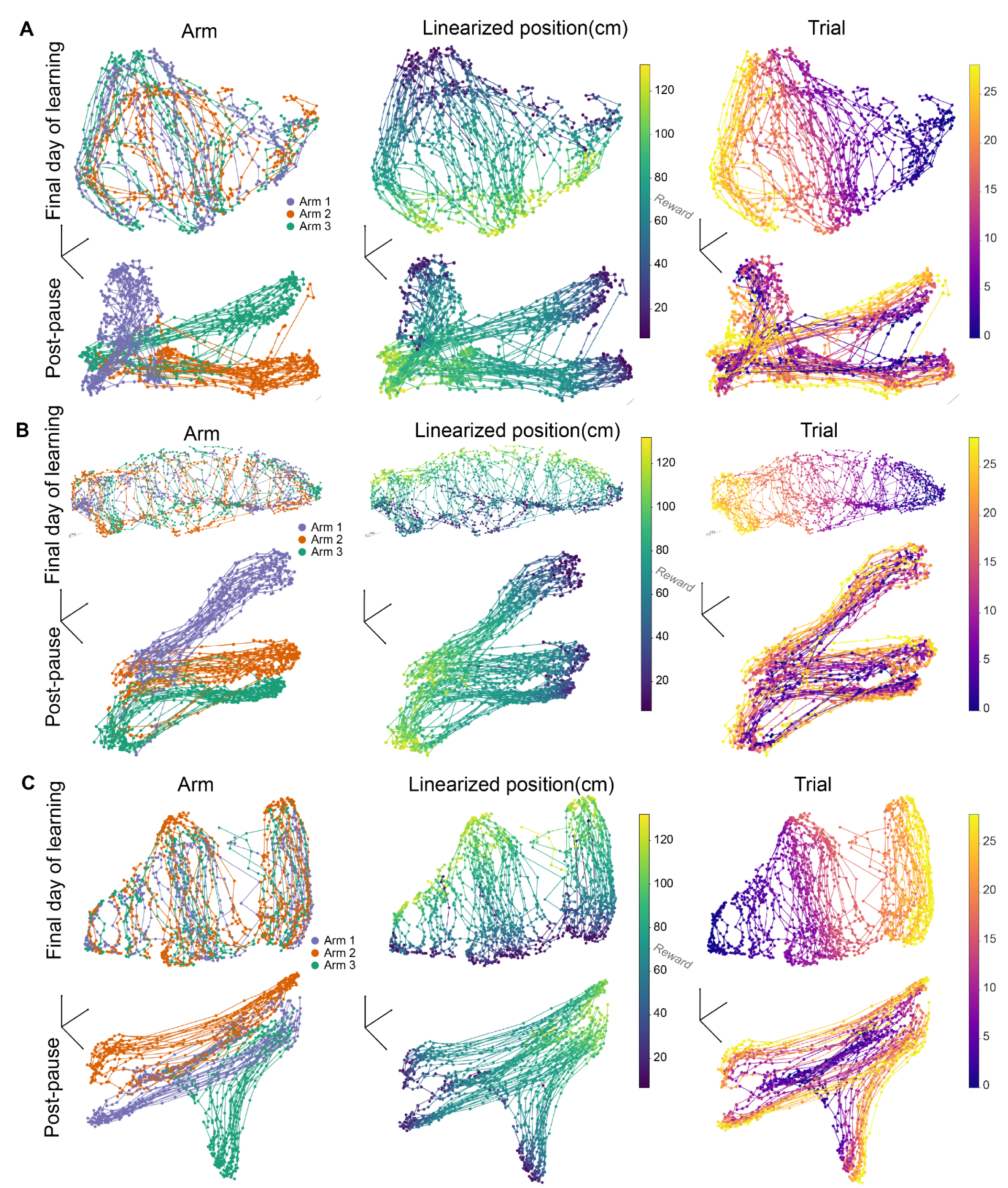
Individual animal examples of mPFC manifold reorganization with task repetition. (**A**–**C**) 3D UMAP projections of mPFC population activity for the remaining three individual animals during the SRW task. Top rows: final learning day. Bottom rows: post-pause day. Manifolds are colored by (left to right): arm identity, linearized position along the entire trajectory, and trial progression.

**Supplementary Figure S9.**
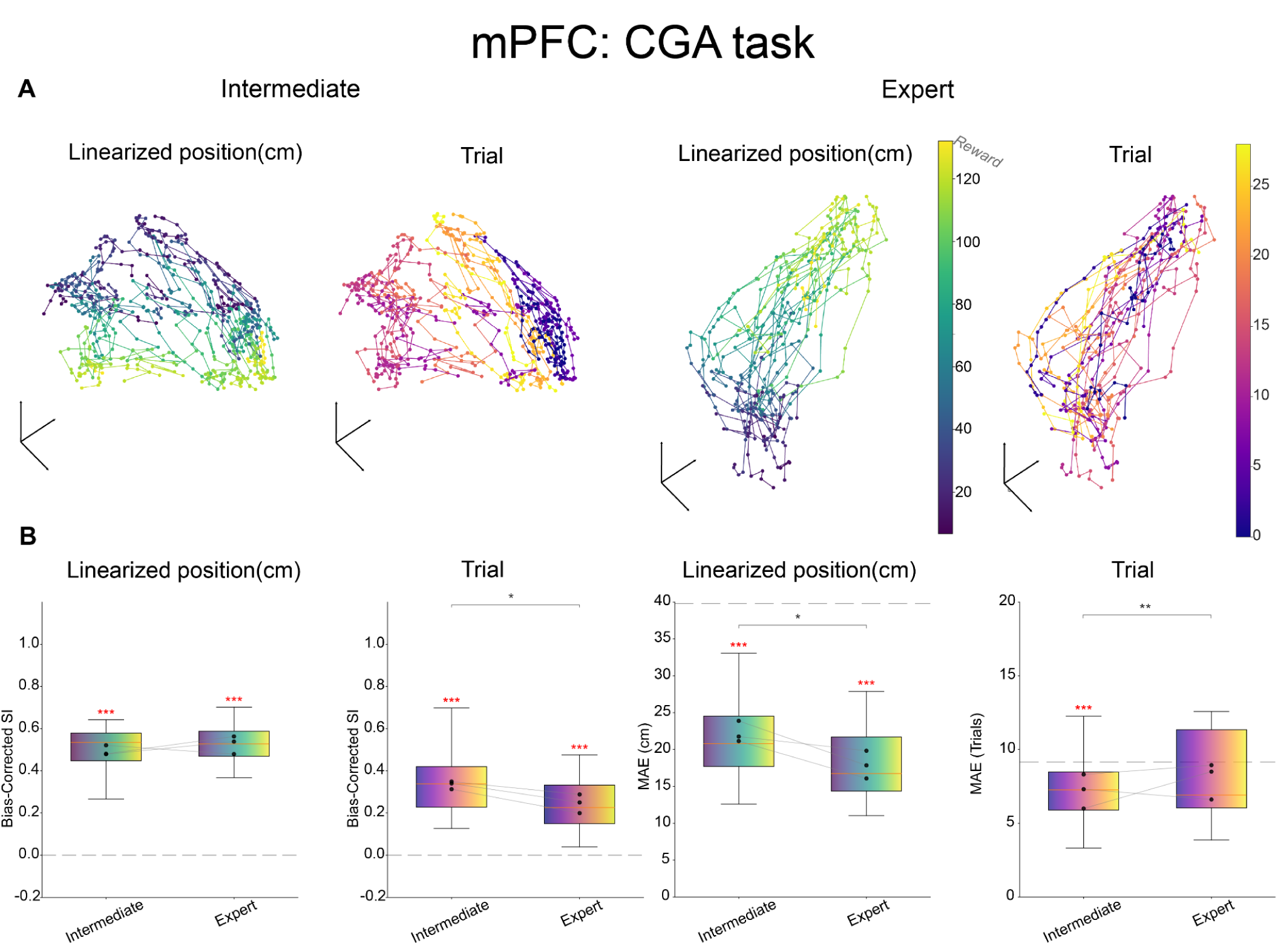
Manifold representation in the mPFC during the CGA task at different learning stages. (**A**) UMAP manifolds before the animals reached the performance criterion (intermediate) (*Right*) and after it was reached (expert) (*Left*). (**B**) Bias-corrected SI and MAE from UMAP manifolds for location and trial progression. Arm identity was not examined because during intermediate learning stages, animals often visited only a single arm, yielding insufficient data for the comparison. Gray lines: medians for individual animals. Dashed lines: chance levels. Red *: significant differences compared to shuffled distributions, Wilcoxon test in (bias corrected SI); Mann-Whitney test in (MAE); black *: significant differences between task variables, * p < 0.05, *** p < 0.001; Wilcoxon test with Holm-Bonferroni correction. differences between task variables

**Supplementary Figure S10.**
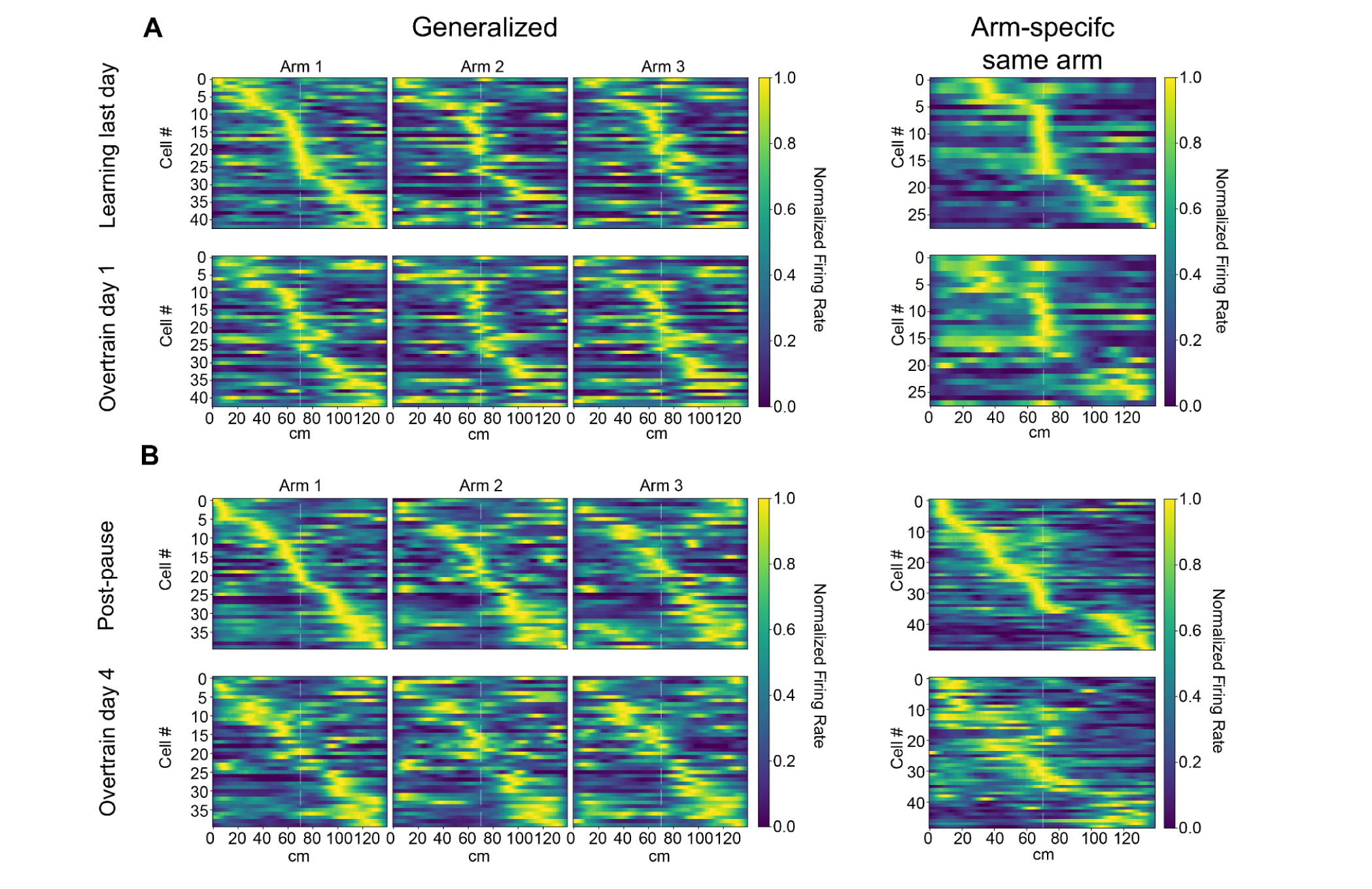
Longitudinal tracking of mPFC spatial rate maps across learning days. (**A**, **B**) Normalized spatial firing rate heatmaps of individual, longitudinally tracked mPFC neurons (see Figure 4C). Tuning transitions between early consolidation (**A**; final learning day vs. first overtraining day) and late consolidation (**B**; overtraining day 4 vs. post-pause day). Each row within a column represents the same cell tracked across the respective days.

**Supplementary Figure S11.**
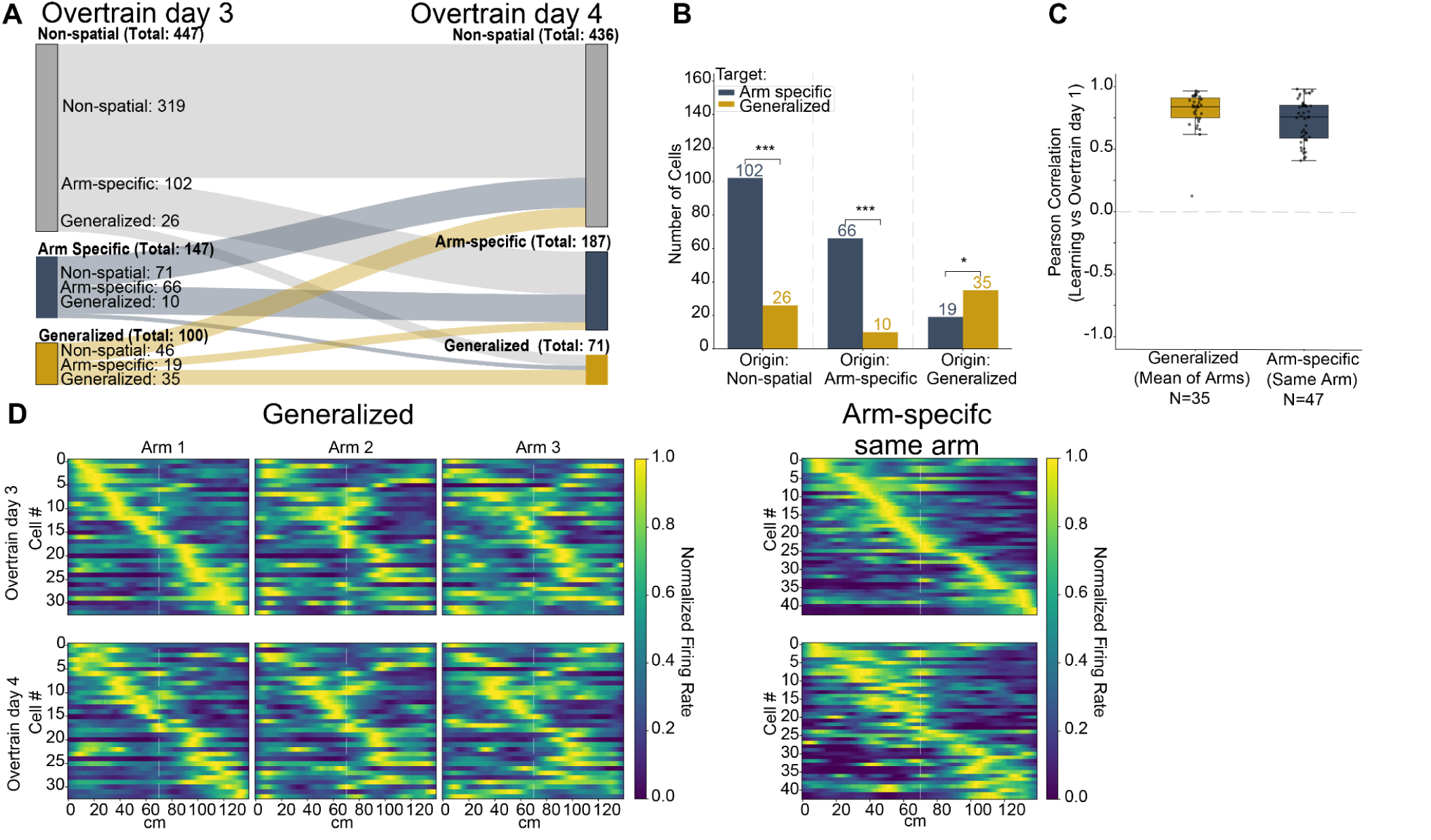
Longitudinal tracking and dynamic routing of mPFC spatial representations during overtraining. (**A**) Sankey diagram illustrating the transition of individual mPFC neurons between coding categories (Non-spatial, Arm-Specific, and Generalized) from overtraining day 3 to overtraining day 4. (**B**) Quantification of cell coding changes. Bars: number of cells transitioning into Arm-Specific or Generalized representations, grouped by their original coding state (* p < 0.05, *** p < 0.001; binomial test). (**C**) Pearson correlations of spatial rate maps for identical cells tracked across the two overtraining sessions. Arm-specific neurons are categorized as “Same Arm” if they maintained their coding on the same arm (47 out of the 66 neurons). (**D**) Normalized spatial firing rate heatmaps of individual, longitudinally tracked mPFC neurons across overtraining day 3 (top rows) and overtraining day 4 (bottom rows). Columns: representative examples corresponding to the Generalized, Arm-specific (same arm) coding categories.

**Supplementary Figure S12.**
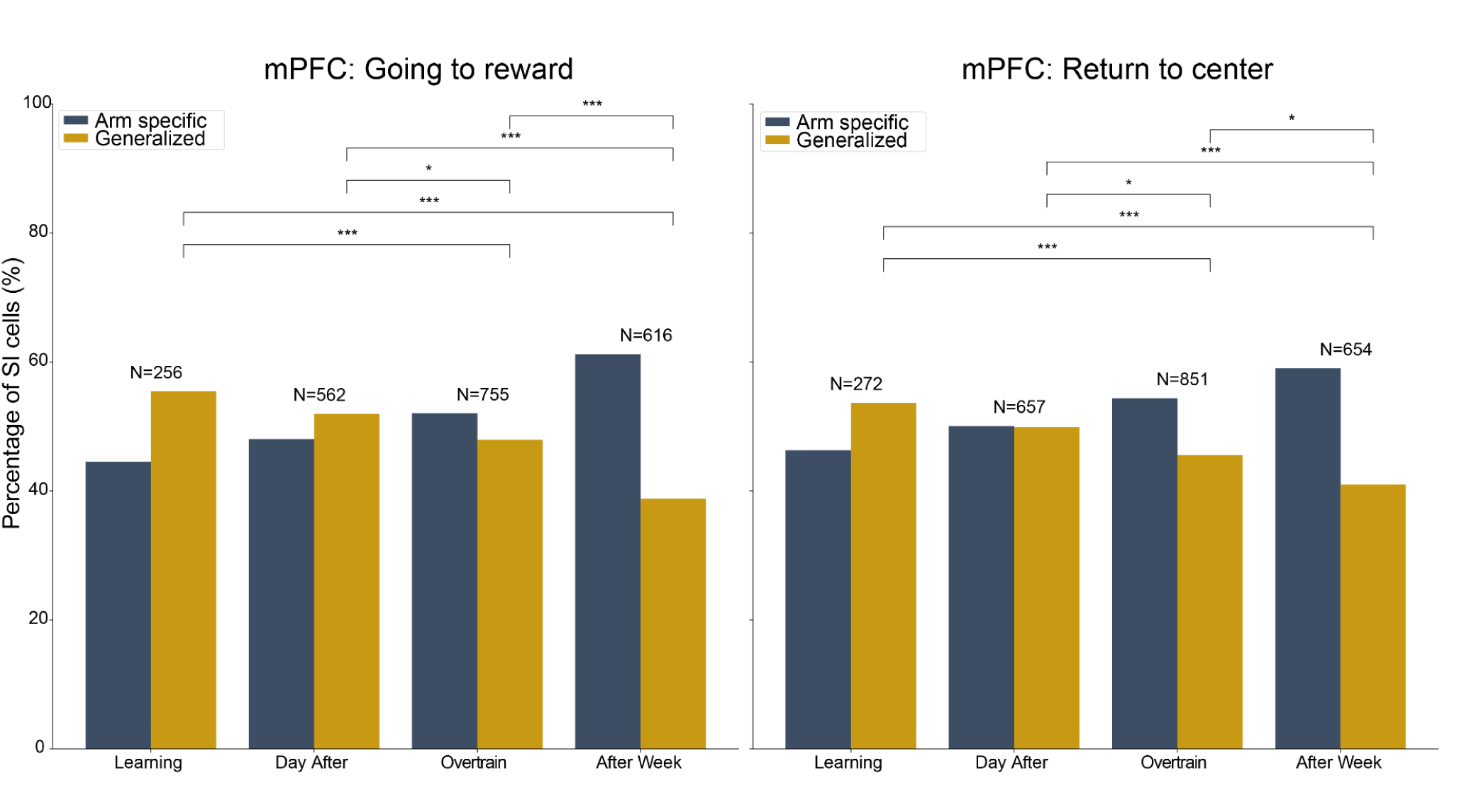
Arm-specific and generalized coding cells in CA1 and mPFC across learning stages in the SRW task. Proportions of arm-specific (blue) and generalized (mustard yellow) neurons in the mPFC during outbound (center to reward; left) and inbound (return to center; right) trajectories. Data across four stages: final learning day, first overtraining day (Overtraining: day 1), subsequent days before the break (Overtraining: rest), and post-pause day. The total number of spatial information-filtered neurons is shown for each category. A progressive increase in arm-specific representations in mPFC occurred as animals reached the overtrained stage. * p < 0.05, p < 0.01, *** p < 0.001 binomial test.

**Supplementary Figure S13.**
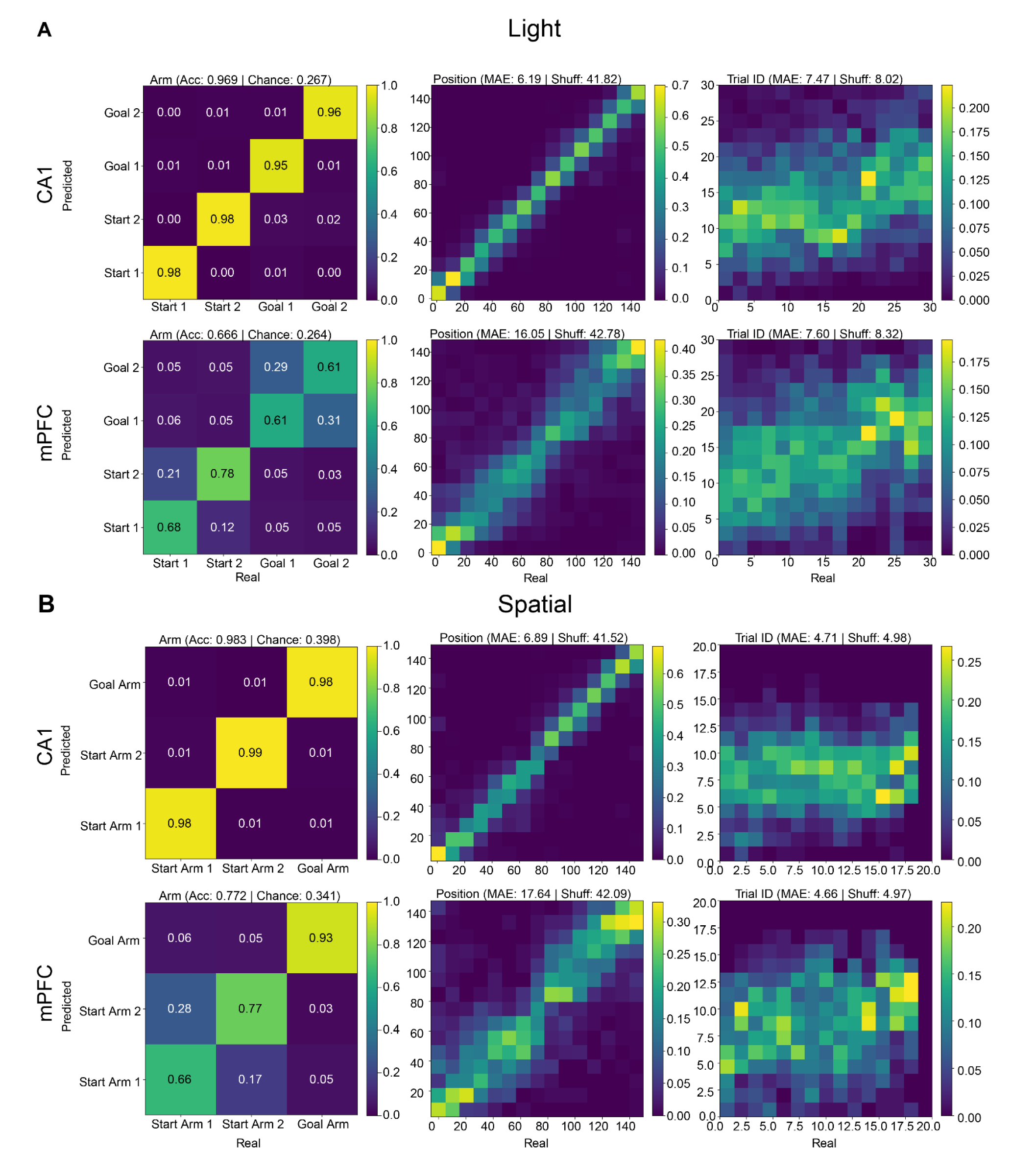
Detailed UMAP decoding performance during Light and Spatial rule tasks. (**A**, **B**) Detailed UMAP-based decoding results for CA1 and mPFC populations during the Light Rule (**A**) and Spatial Rule (**B**) paradigms (see Figure. 5). Left to right: confusion matrices for arm identity (evaluating start and goal arms), and joint distributions for predicted versus real linearized position and trial progression (Trial ID). Values above plots: overall model accuracy (arm identity) or mean absolute error (position and trial ID), alongside chance levels calculated from shuffled distributions. Data were pooled together across all animals

**Supplementary Figure S14.**
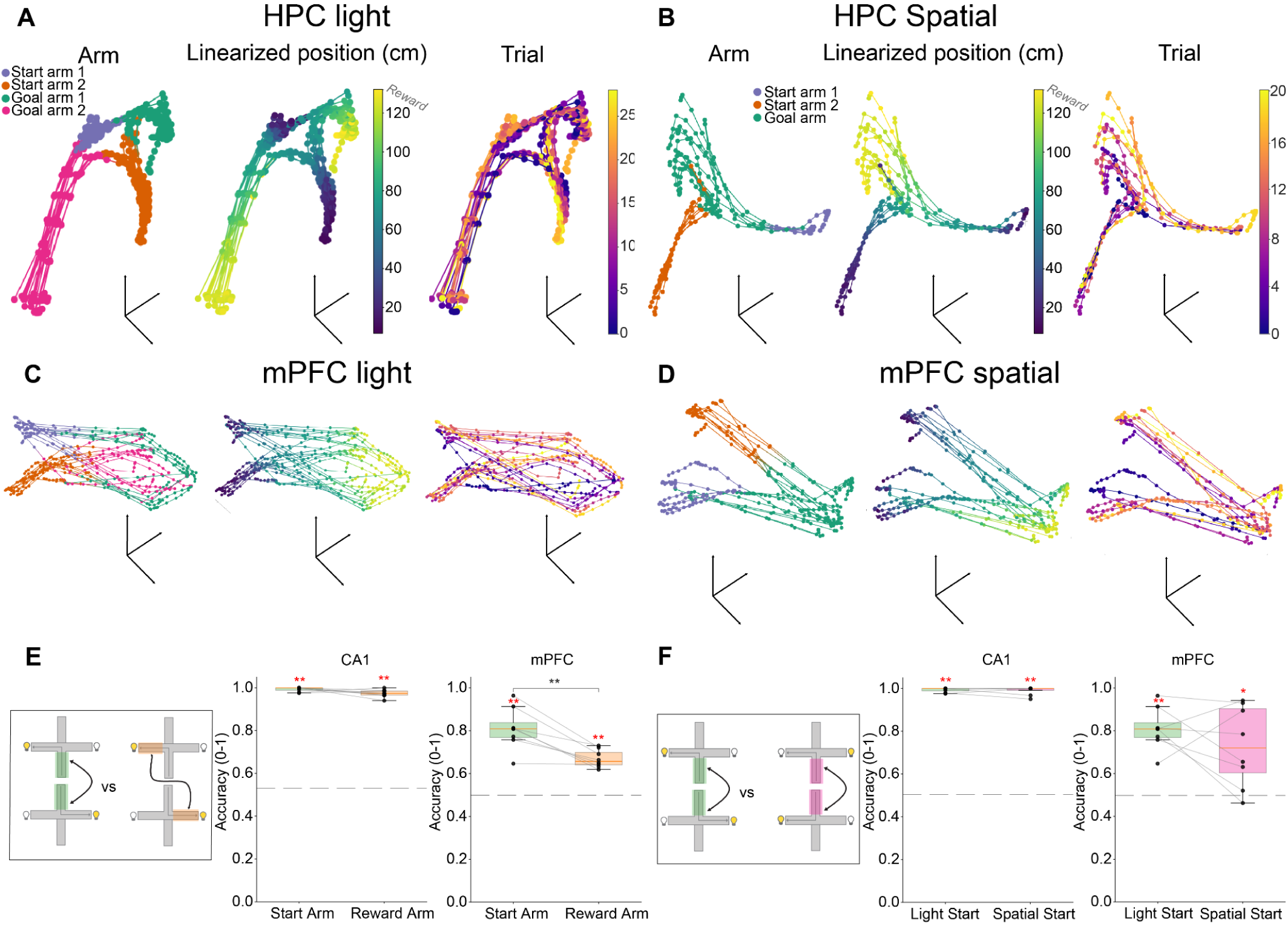
UMAP manifold examples and decoding performance across navigation rules. (**A**–**D**) Individual session 3D UMAP projections for CA1 (**A**, **B**) and mPFC (**C**, **D**) populations during Light and Spatial rules. Manifolds are colored by (left to right): arm identity (encompassing start and goal arms), linearized position, and trial progression. (**E**) UMAP-based decoding accuracy comparing the Start Arm versus Reward Arm exclusively during the Light Rule. (**F**) Comparison of Start Arm decoding accuracy between the Light Rule and Spatial Rule paradigms. Gray lines: individual recording sessions. Dashed lines: chance levels. Red *: significant differences compared to shuffled distributions, Mann-Whitney test; black *: significant differences between task variables or conditions, * p < 0.05, p < 0.01; Wilcoxon test with Holm-Bonferroni correction.

